# Targeting Rho Kinase in Proliferative Vitreoretinopathy with Netarsudil: Multimodal Preclinical Evidence and Exploratory Clinical Application

**DOI:** 10.64898/2026.11.20.768165

**Authors:** Jeysson Sanchez-Suarez, Bernardo Bach, William P. Miller, Anil Upreti, Nakyung Ryoo, Rose Lin, Dhanesh Amarnani, Daisy Y. Shu, Aruvi Vijikumar, Karim W. Barake, Yvonne Adu-Rutledge, Max Semegran, Bryan A. Kaplan, Audrey L. Gunawan, Jonathan F. Russell, Romy P. Bejjani, Kunho Bae, Padmanabhan Pattabiraman, Anton Lennikov, Elizabeth J. Rossin, Dean Eliott, Edward F Linton, Elliott H Sohn, Michael O’Hare, Leo A. Kim

## Abstract

**Objective:** To investigate the involvement of Rho-kinase signaling in proliferative vitreoretinopathy-associated fibrosis and to evaluate the effects of pharmacologic Rho-kinase inhibition with netarsudil across experimental models and clinical outcomes.

**Design:** Experimental study with in vitro and ex vivo testing in patient-derived models, in vivo evaluation in a rabbit model, and analysis of clinical outcomes in patients with proliferative vitreoretinopathy.

**Samples and Subjects:** Patient-derived primary proliferative vitreoretinopathy cells (C-PVR cells), ex vivo proliferative vitreoretinopathy membrane explants, a rabbit model of proliferative vitreoretinopathy, and patients with grade C proliferative vitreoretinopathy, including a prospective treatment cohort and retrospective controls.

**Methods:** Single-cell RNA sequencing was used to assess Rho-kinase pathway activation in human proliferative vitreoretinopathy membranes. The effects of the Rho-kinase inhibitor netarsudil were evaluated in C-PVR cells using in vitro assays of cytotoxicity, proliferation, cellular migration, and contraction; in ex vivo proliferative vitreoretinopathy membrane explants; in an in vivo rabbit model of proliferative vitreoretinopathy; and through repeated event analysis of clinical outcomes following topical netarsudil administration.

**Main Outcome Measures:** Rho-kinase pathway activation, cytotoxicity and cellular proliferation, cellular migration and contraction, disease severity in an animal model of proliferative vitreoretinopathy, and surgical reoperation rates in patients.

**Results:** Rho-kinase pathway signaling was enriched in fibrotic cell populations within human proliferative vitreoretinopathy membranes. In C-PVR cells, netarsudil exhibited concentration-dependent effects, with cytotoxicity observed at concentrations of 5 µM and higher, whereas at 2.5 µM netarsudil was not cytotoxic and preserved antiproliferative activity while significantly inhibiting cellular migration and contraction. Netarsudil also reduced fibrotic remodeling in ex vivo membrane explants and attenuated disease severity in a rabbit model. In patients with grade C proliferative vitreoretinopathy, required zero reoperations, compared to a 40% rate in untreated eyes from the retrospective cohort (HR = 0.097 [0.0008-0.75], p = 0.02).

**Conclusions:** Rho-kinase signaling is implicated in fibrotic remodeling associated with proliferative vitreoretinopathy. Pharmacologic Rho-kinase inhibition with netarsudil attenuated fibrotic phenotypes across experimental models in a concentration-dependent manner, and topical administration was associated with low reoperation rates, supporting further investigation.

## Introduction

Proliferative vitreoretinopathy (PVR) is the principal cause of surgical failure following retinal detachment (RD) repair, with no FDA-approved treatments available ^1^. Despite decades of investigation, current interventions remain surgical, and re-detachment of the retina due to contractile fibrotic membrane formation often results in irreversible vision loss ^2^. Although extensively studied, pharmacological treatments for PVR are still lacking ^3^, leaving repeated retinal surgery to remove fibrotic membranes as the only option.

Surgical success hinges on the complete removal of membranes, pathologic cells, and vitreous cortex remnants. Residual tissue and cells can increase recurrence rates, compromising surgical outcomes ^4^. PVR contributes to retinal re-detachment in approximately 10-15% of cases following rhegmatogenous RD (RRD) surgery ^5^ and affects 40-60% of individuals with severe ocular trauma ^6^. In such cases, reattachment rates are significantly lower, often requiring multiple surgeries and leading to poorer functional outcomes ^7^. The pressing need for alternative therapeutic options highlights an important clinical gap.

The pathogenesis of PVR is complex and multifactorial, involving various cell types-including retinal pigment epithelial (RPE) cells, glial cells, and macrophages--and pro-inflammatory and pro-fibrotic cytokines ^3^. Among these, transforming growth factor-beta 2 (TGF-β2) plays a pivotal role in driving PVR pathogenesis ^8^. Elevated TGF-β2 levels are associated with epithelial-mesenchymal transition (EMT), increased extracellular matrix (ECM) protein synthesis, and the upregulation of α-smooth muscle actin (α-SMA) expression, all of which are associated with myofibroblast transdifferentiation ^8,9^.

The Rho GTPase-ROCK pathway drives key processes involved with mesenchymal transition, including cell proliferation, migration, contraction, and fibrosis ^10^. By regulating the actin cytoskeleton, ROCK facilitates changes in cell morphology and motility. These changes, characterized by the loss of epithelial markers such as E-cadherin and the upregulation of mesenchymal markers such as vimentin, are hallmarks of EMT ^11^. ROCK activation reorganizes the cytoskeleton, altering cell morphology and behavior, promoting actin stress fiber formation, and enhancing cell motility and contractility ^12^. Notably, TGF-β and ROCK signaling are interconnected, as TGF-β activates ROCK, which in turn amplifies TGF-β signaling ^13^.

We hypothesized that pharmacologic inhibition of the ROCK signaling pathway would mitigate key fibrotic behaviors in PVR and reduce surgical failure. Here, we combined single-cell transcriptomic profiling of patient-derived membranes with functional validation *in vitro*, *ex vivo*, and in a rabbit model, as well as a pilot clinical study of patients with grade C PVR treated postoperatively with topical netarsudil, a potent inhibitor of ROCK1 and ROCK2^14^. Our study demonstrates that a topically administered drug can prevent the need for reoperation in severe PVR. This work highlights the translational potential of repurposing a widely available, FDA-approved agent to address an urgent unmet need in retinal surgery.

## RESULTS

### ROCK pathway genes are expressed in patient-derived PVR membranes

To characterize transcriptional heterogeneity and ROCK pathway gene expression in PVR membranes, we evaluated a previously generated single-cell RNA sequencing (scRNA-seq) dataset from our lab comprising patient-derived PVR membranes from eight donors **(Fig. 1A)** ^15^. Our revised analysis identified 13 transcriptional clusters **(Fig. 1B).** Based on their transcriptomic profiles, clusters were assigned reference annotations using scType ^16^ as follows: macrophage, fibroblast, natural killer/T-cell, astrocyte, retinal pigment epithelial (RPE), monocyte, myeloid dendritic, amacrine, B-cell, retinal progenitor, pericyte, endothelial, and cone bipolar. These labels were used as computational descriptors of marker-based transcriptional similarity, rather than definitive evidence of cell identity or tissue of origin. Six scType-annotated groups-macrophage, fibroblast, natural killer/T-cell, astrocyte, RPE, and monocyte-accounted for approximately 92% of the analyzed cells **(Fig. 1C).** Previous reports have also described cellular heterogeneity in PVR membranes ^17,18^.

**Figure 1.**
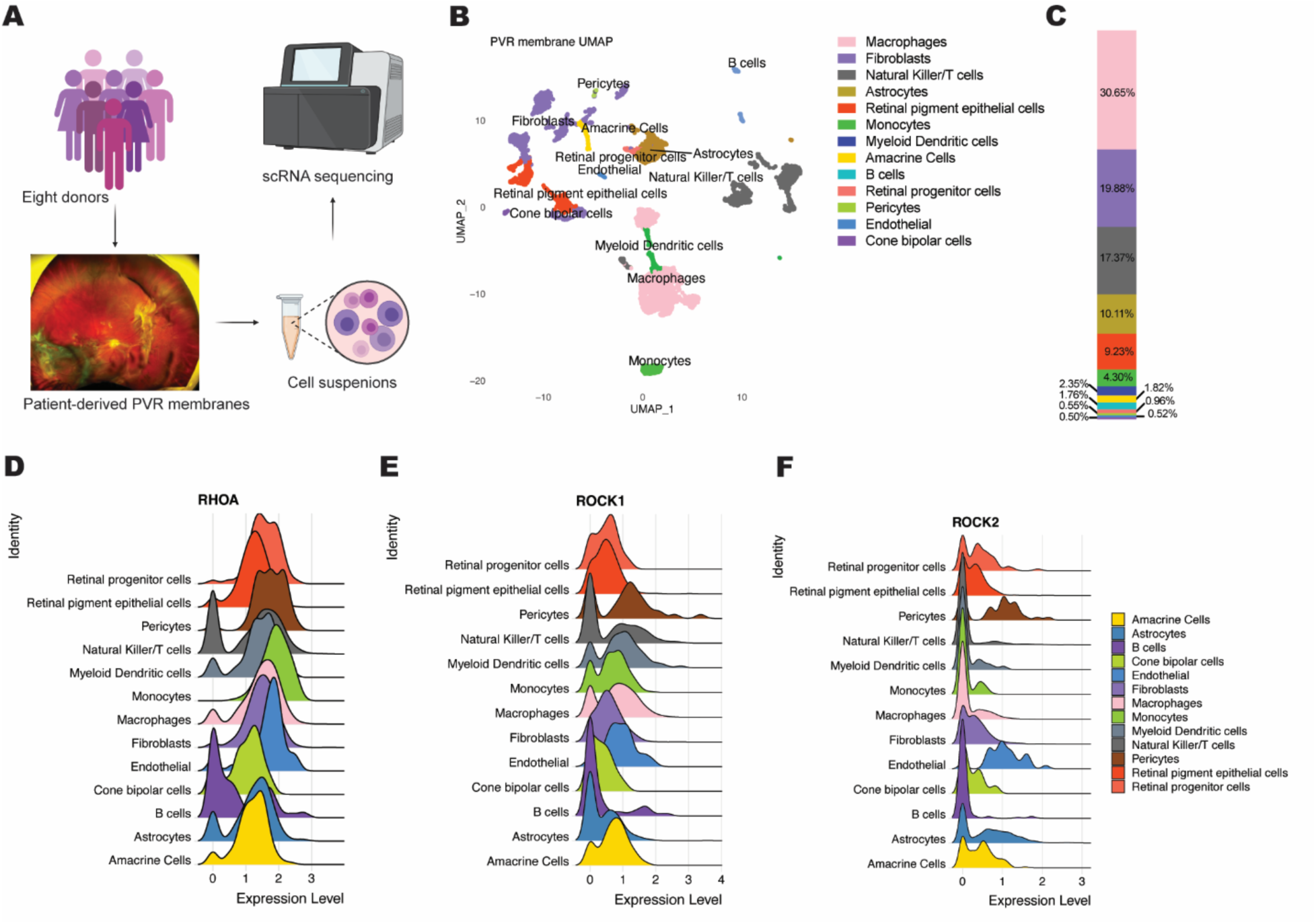
Single-cell characterization of Rho kinase pathway in patient-derived PVR membranes. **(A)** Workflow diagram for scRNA-seq analysis of PVR patient membrane samples (n=8). **(B)** UMAP dimensionality reduction was performed to visualize 13 different cell clusters. **(C)** Stacked bar charts showing the cell composition in PVR membranes. **(D-F)** Ridgeline plots showing relative expression of **(D)** RhoA **(E)** ROCK1 **(F)** ROCK2 across each cell type.

Given that the ROCK pathway has previously been implicated in EMT and hypothesized as a mediator of PVR pathogenesis ^19^, we evaluated the expression profiles of RHOA, ROCK1, and ROCK2 across the scType-annotated groups. RHOA transcripts were detected across multiple annotated groups (**Fig.** 1D). ROCK1 expression was also evident in multiple groups, including those annotated as fibroblast, macrophage, monocyte, and natural killer/T-cell **(Fig. 1E)**. ROCK2 expression varied across the annotated groups, with expression evident in groups annotated as astrocyte, endothelial, and pericyte **(Fig. 1F)**. Together, these findings document the expression of RHOA-ROCK pathway components within the transcriptionally heterogeneous PVR membrane dataset.activity or isoform-specific functional roles in PVR.

### Netarsudil inhibits PVR-associated cellular responses

-We evaluated netarsudil in primary cultures derived from patient PVR membranes (C-PVR cells). LDH release increased at tested concentrations of 5 µM or higher, whereas 2.5 µM did not produce a detectable increase at the assessed time points **(Fig. 2A)**. Complementary assays showed increased Annexin V binding and loss of membrane integrity with 5 and 10 µM netarsudil, but no corresponding increase with 2.5 µM **(Fig. 2B-C)**. The earlier appearance of Annexin V signals relative to membrane-integrity signals was consistent with apoptotic events preceding membrane disruption at the higher concentrations. These results supported selection of 2.5 µM for subsequent experiments to obtain robust biological activity without detectable cytotoxicity in these assays under the tested conditions.

**Figure 2.**
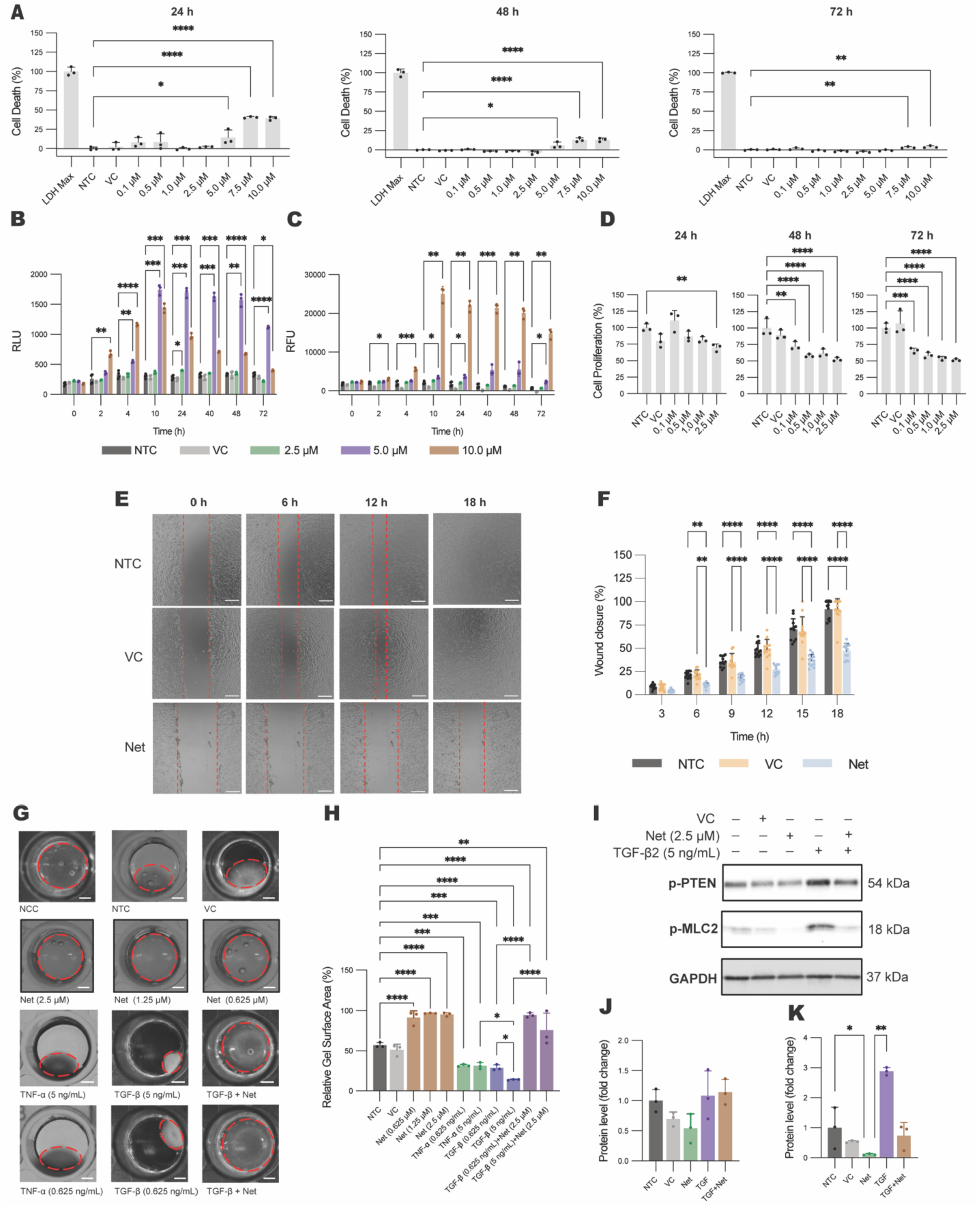
Functional effect of Rho Kinase inhibition in PVR cells. **(A)** Bar graph showing the percentage of cell death at different concentrations (0.1 to 10 µM) of netarsudil, vehicle control (VC), and non-treated cells (NTC) at 24, 48 and 72 hours of exposure; (n=3). **(B)** Bar graph showing the kinetics of the annexing V binding signal (measured in relative luminescence units, RLU) at different concentrations (2.5, 5.0, and 10 µM) of netarsudil, VC, and NTC over 72 hours; (n=3). **(C)** Bar graph showing the kinetics of the loss of membrane integrity signal (measured in relative fluorescence units, RDU) at different concentrations (2.5, 5.0, and 10 µM) of netarsudil, VC, and NTC over 72 hours; (n=3). **(D)** Bar graph showing the percentage of cell proliferation at different concentrations (0.1 to 2.5 µM) of netarsudil, VC, NTC at 24, 48 and 72 hours of exposure. **(E)** Representative time-lapse (0, 6, 12, and 18 hours) images of wound healing closure; scale bar=200 μm. **(F)** Graph showing the time course of the wound area in the wells treated with netarsudil, VC, and NTC.; (n=12) **(G)** Representative images of collagen matrixes; scale bar=2 mm. **(H)** Bar graph showing the relative gel surface area (in percentage) after 72 hours of treatments, i.e., collagen without cells (non-cell control NCC), NTC, VC, treated with netarsudil at 0.635, 1.25 and 2.5 µM, TNF-α at 0.635 and 5 ng/mL, TGF-β2 at 0.625 and 5 ng/mL, TGF-β2 at 0.625 ng/mL plus netarsudil at 2.5 µM and TGF-b at 5 ng/mL plus netarsudil at 2.5 µM; (n=3). **(I)** Representative immunoblot showing reduced phosphorylation of PTEN and MLC2 after netarsudil treatment. **(J and K)** Bar graph showing the quantification of immunoblot (as shown in **i**) results for phospho-PTEN **(J)** and phospho-MLC2 **(K)**; (n=3). (n =) indicates total technical replicates across 3 independent experiments. Data are represented as mean ± S.E.M. *P < 0.05, **P<0.01, ***P<0.001.

Netarsudil reduced C-PVR cell proliferation, with significant inhibition at the higher tested concentrations after 48 and 72 hours **(Fig. 2D)**. At 2.5 µM, relative proliferation was 52.4% and 51.4% of untreated controls at 48 and 72 hours, respectively. In the scratch-wound assay, netarsudil-treated cultures showed less wound closure, reaching 47.1% at 18 hours compared with 92.1% and 90.9% in untreated and vehicle-treated controls, respectively **(Fig. 2E-F)**.

Netarsudil also markedly inhibited collagen gel contraction **(Fig. 2G-H)**. Relative gel surface area was 91.22 ± 8.71%, 96.85 ± 0.55%, and 95.30 ± 2.39% with 0.625, 1.25, and 2.5 µM netarsudil, respectively, compared with 51.16 ± 7.00% in vehicle-treated controls. At 0.625 and 5 ng/mL TGF-β2, relative gel surface area increased from 28.91 ± 3.69% and 14.01 ± 0.68% without netarsudil to 94.45 ± 2.90% and 75.66 ± 21.38%, respectively, with 2.5 µM netarsudil, indicating attenuation of TGF-β2-associated contraction.

We next examined the effects of netarsudil on p-PTEN and p-MLC2 levels by Western blotting **(Fig. 2I-K)**. No statistically significant differences in p-PTEN levels were detected among treatment groups. Netarsudil alone significantly reduced p-MLC2 levels compared with untreated controls, consistent with the observed inhibition of gel contraction.

### Netarsudil treatment is associated with changes in C-PVR transcriptomic profiles

To characterize the transcriptional changes associated with netarsudil treatment, we performed scRNA-seq of C-PVR cells exposed to vehicle (VC) or 2.5 µM netarsudil for 48 hours **(Fig. 3A and 3B).** The scType-assigned labels were manually reviewed and revised to reflect the closest transcriptional state supported by the observed gene expression profiles, without implying definitive cell identity or origin. The original scType annotations and corresponding revised labels are provided in Supplementary Table X. Netarsudil treatment was associated with a marked redistribution of the annotated transcriptional states. The proportion of analyzed cells assigned the extracellular matrix (ECM) label decreased from 88.6% in VC to 3.3% after treatment, while the fibroblast-annotated group decreased from 2.9% to 1.97%. In contrast, the RPE-annotated group increased from 1.33% to 4.7% **(Fig. 3C)**. The treated cultures were predominantly represented by groups annotated as “Glycinergic amacrine cells” (43.42%) and “Unknown” (43.24%) **(Fig. 3B and 3C)**. These changes indicate a shift in the relative representation of transcriptional profiles within the analyzed cultures; they do not establish selective loss or emergence of defined cell types.

**Figure 3.**
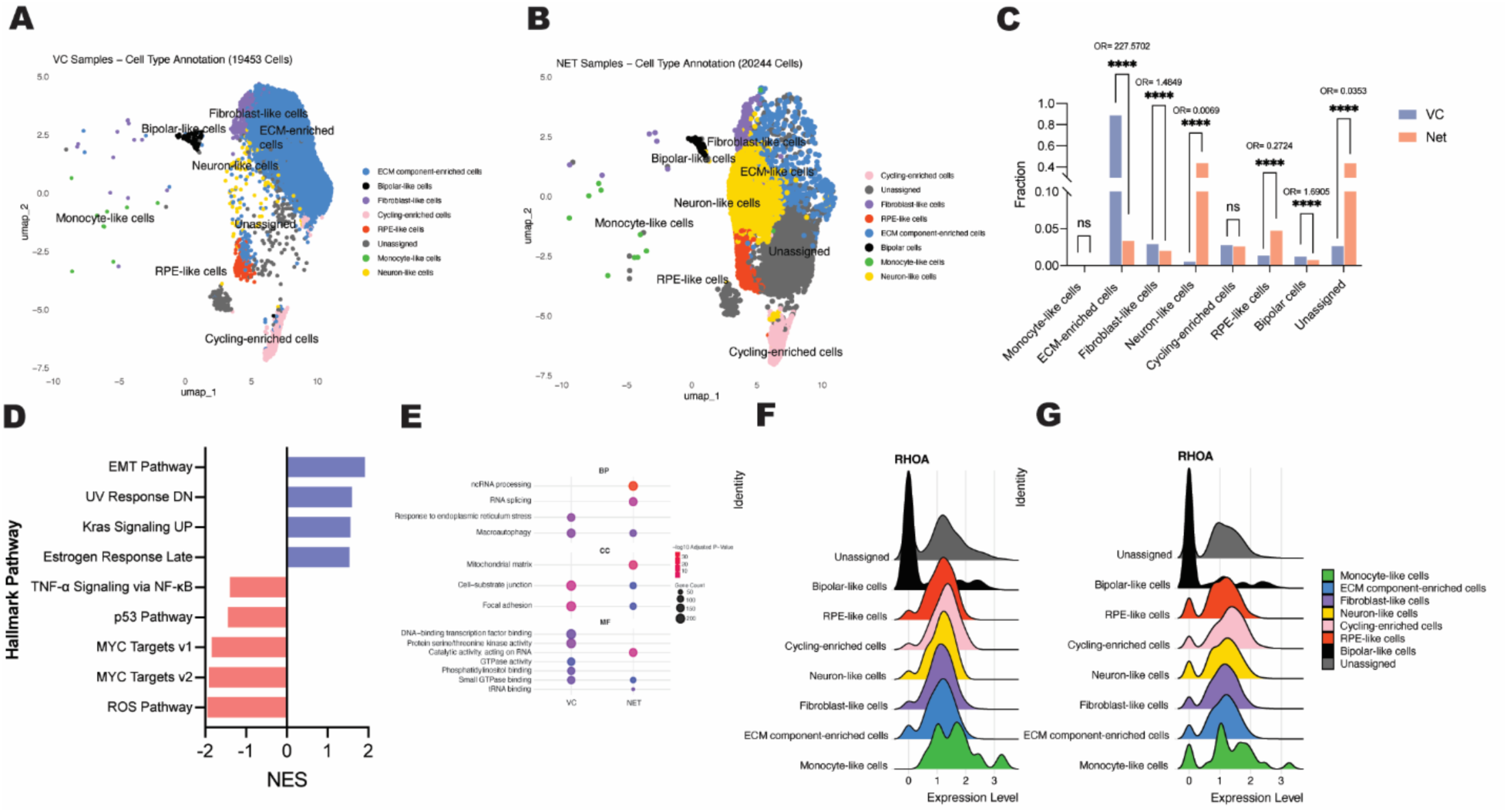
Mechanism of action for netarsudil in C-PVR cells. **(A)** UMAPs of C-PVR cells from vehicle control displaying a combined 8 cell types distinguished by colors (n=2). **(B)** UMAPs of C-PVR cells after netarsudil (2.5µM) treatment for 48 hours (n=2). **(C)** Bar graph showing C-PVR subpopulation in vehicle control and netarsudil (2.5 µM) treated cells at 48 hours. **(D)** SEA analysis showing hallmark pathways enriched in the vehicle compared to netarsudil. **(E)** Gene ontology terms of interest including biological processes (BP), cellular component (CC), and molecular function (MF). **(F and G)** RhoA expression displayed as a ridge plot for **(F)** VC and **(G)** (n= two independent cultures) netarsudil-treated groups. ****P<0.0001 vs VC.

Gene set enrichment analysis (GSEA) showed that the shift away from ECM-annotated profiles was accompanied by differences in fibrosis-related gene expression. EMT was the most enriched Hallmark gene set in VC relative to netarsudil, consistent with a lower EMT-associated transcriptional signature after treatment **(Fig. 3D)**. UV Response DN, KRAS Signaling UP, and Estrogen Response Late were also enriched in VC. Conversely, TNF-α Signaling via NF-κB, p53 Pathway, MYC Targets V1/V2, and Reactive Oxygen Species Pathway were enriched after netarsudil treatment. Thus, the lower EMT-associated signature was accompanied by stress-related transcriptional responses. At the gene level, SNAI2, an EMT-associated transcriptional regulator ^20^, and the matrix-associated genes *COL5A2*, *COL3A1*, *MMP3*, and *TIMP3* showed higher expression in VC than in the netarsudil group **(Table S1)**. Additional expression differences involved genes related to proliferation, migration, and contractility, providing transcriptional context for the changes observed in the functional assays. Gene-level results and average log2 fold changes are presented in Table S1.

Gene ontology (GO) analysis further characterized the treatment-associated expression differences. VC-associated terms included endoplasmic reticulum stress response and macroautophagy (biological processes), focal adhesion and cell-substrate junctions (cellular components), and phosphatidylinositol binding, small GTPase binding, protein serine/threonine kinase activity, and DNA-binding transcription factor binding (molecular functions). Netarsudil-associated terms included RNA splicing and ncRNA processing, mitochondrial matrix, tRNA binding, and catalytic activity acting on RNA **(Fig. 3E)**. These results extend the transcriptional changes beyond EMT to adhesion-associated and RNA-processing functions. We also used CellChat to explore predicted ligand-receptor signaling among the annotated groups ^19^. In the RPE-annotated group, predicted incoming pathways included IGF, VEGF, PARs, GRN, GAS, PROS, NT, and PSAP in VC, whereas no incoming pathways were inferred after treatment under the applied analysis settings. Predicted outgoing pathways differed between conditions: BMP, WNT, GAS, and NT in VC, compared with GAS, NRG, and WNT after netarsudil treatment **(Fig. S1)**. These expression-based predictions suggest changes in potential signaling relationships and require functional validation.

Finally, RHOA expression distributions shifted toward more cells with undetected transcripts in several annotated groups after netarsudil treatment **(Fig. 3F and 3G).** Exploratory comparisons yielded reported p-values of 0.0162 for the RPE-annotated group, 0.0473 for the rod bipolar-annotated group, and 0.0002 for the “Unknown” group **(Fig. S2)**. Together, these observations associate netarsudil exposure with changes in transcriptional-state representation, lower EMT-associated gene expression, and altered RHOA transcript distributions.

### Netarsudil suppresses outgrowth of patient-derived PVR explants

We next evaluated netarsudil in explants derived from patient PVR membranes. Explant growth was similar between groups early in culture, with greater outgrowth in vehicle-treated explants after day 8 **(Fig. 4A-B)**. At day 12, growth was significantly lower with 2.5 µM netarsudil than with vehicle (25.7 ± 9.3% vs 100. 0 ± 13.0%; p = 0.0097; **Fig. 4C)**. These findings extend the cell-culture observations to an ex vivo tissue model.

**Figure 4.**
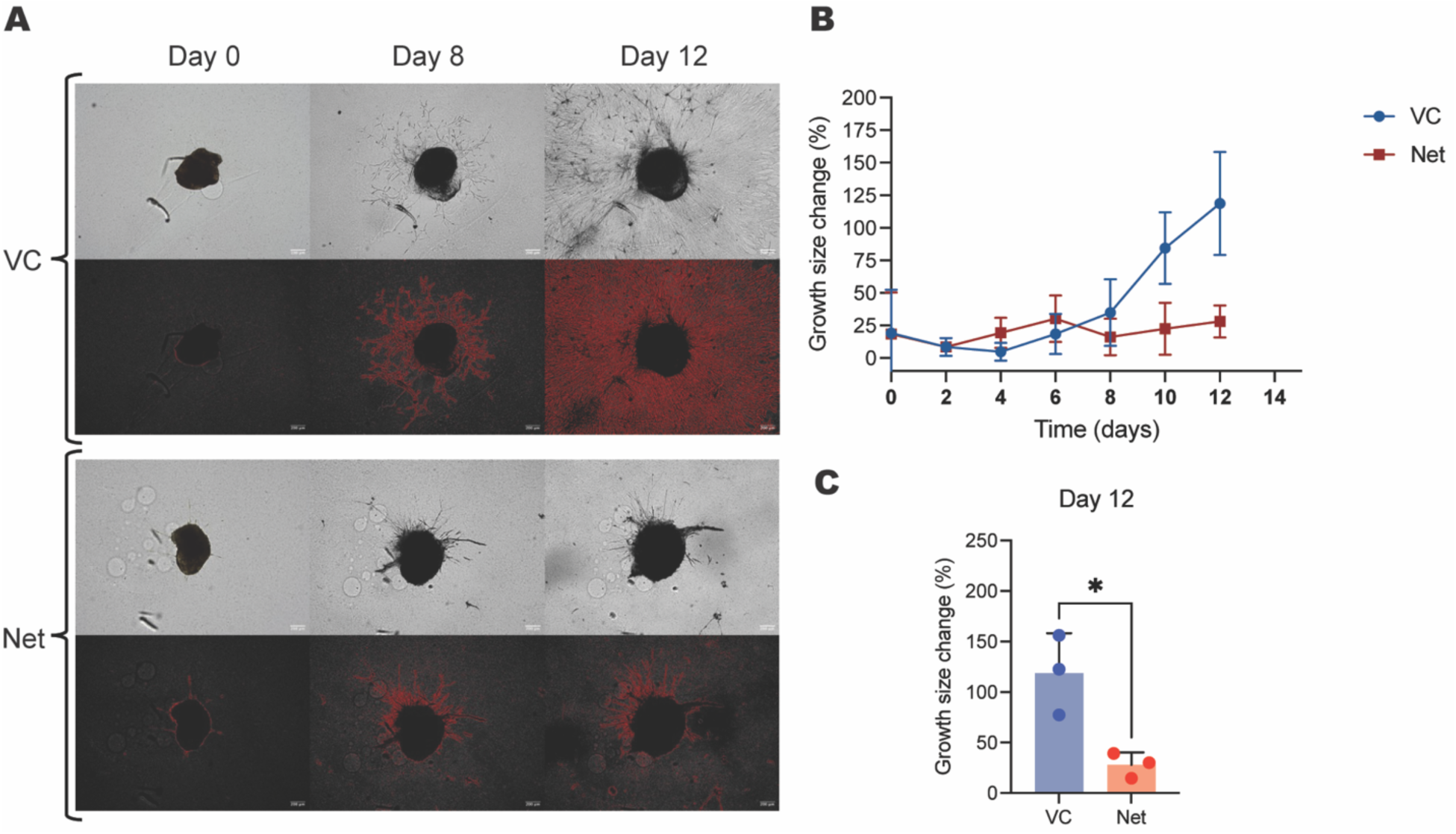
Netarsudil inhibits the growth of PVR patient-derived explants. **(A)** Representative micrographs of explants treated with DMSO (VC) (n=3) or netarsudil (Net) (n=3) at 2.5 μM at 0, 8, and 12 days after treatment. The upper panels in each group present the original explant photograph, while the lower panels utilize red highlighting to delineate the extent of the explant. **(B)** bar graph showing the quantitative analysis of explant growth. The percentage growth mean ± SEM of each explant, corrected against its initial size at the starting point of the treatment period, is plotted. (n=number of technical replicates derived from two patients); *p < 0.05.

### Intravitreal netarsudil attenuates PVR in a rabbit model

Intravitreal administration of Netarsudil attenuated proliferative membrane formation and mitigated associated retinal pathology in a rabbit model of proliferative vitreoretinopathy (PVR).

Two weeks after intravitreal injection of C-PVR cells, OCT and infrared (IR) imaging revealed dense post-lenticular membranes within the anterior vitreous, high cellularity within the vitreous cavity, and extensive epiretinal membranes exerting broad traction on the retina in the vehicle control (VC) group **(Fig. 5A-5D)**. In contrast, eyes treated with Netarsudil exhibited no retrolental fibrotic occlusion and lacked signs of extensive epiretinal proliferation **(Fig. 5E and 5F)**. Histological analysis corroborated the in-vivo findings. H&E staining of VC eyes demonstrated prominent fibrous membranes and retinal distortion consistent with PVR **(Fig. 5G)**. In Netarsudil-treated eyes, the retinal architecture remained largely intact, with minimal membrane formation or retinal disruption **(Fig. 5H)**.

**Figure 5.**
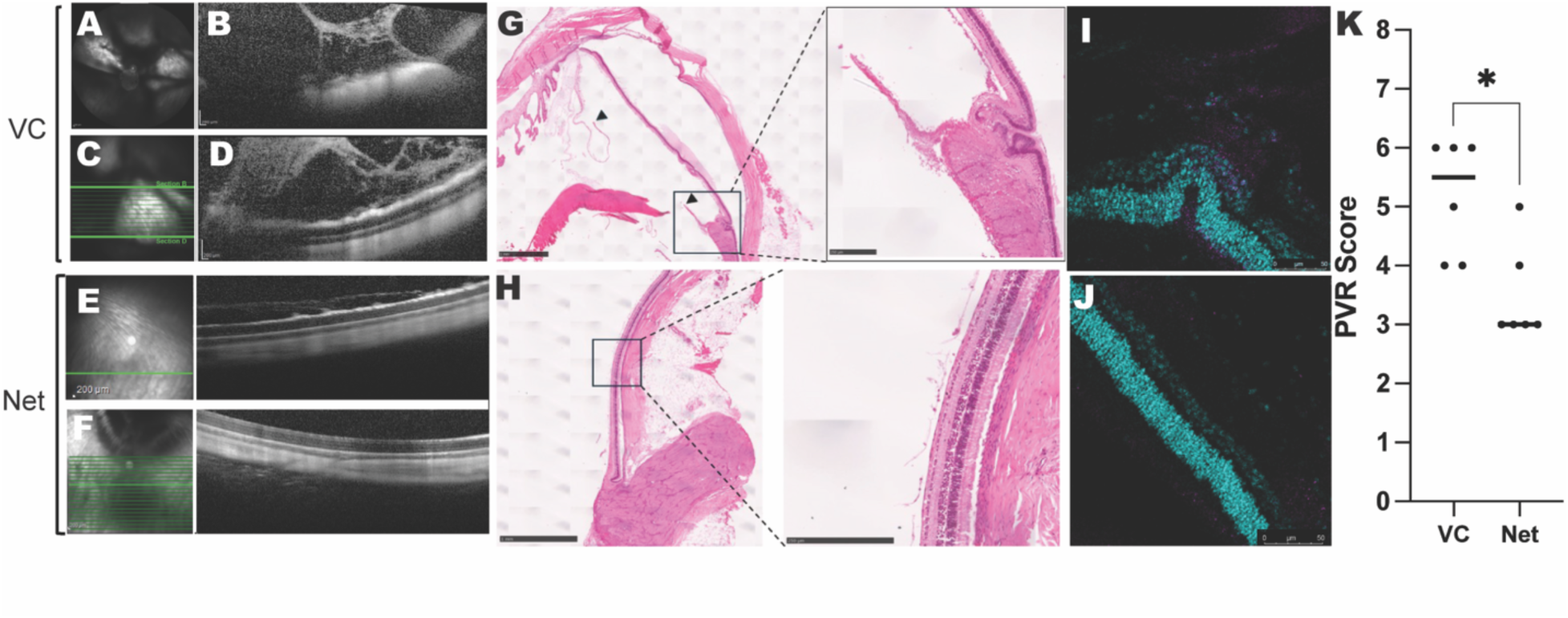
Netarsudil treatment attenuates PVR progression in a rabbit model of PVR. (A-F) Representative infrared (IR) and optical coherence tomography (OCT) images of vehicle control (VC; A-D) and netarsudil-treated eyes. **(A)** IR image showing dense post-lentic membrane formation. **(B, D)** Cross-sectional OCT scans reveal dense epiretinal membranes and intravitreal cellular components exerting traction on the retina (scan lines indicated in **(C))**. **(E)** IR image and OCT scan of Net-treated eyes showing minimal membrane formation in the peripheral retina. **(F)** Quiescent retina in the peripapillary area (inferior to the optic nerve) with no visible proliferative activity. **(G, H)** Histological sections of PVR-induced eyes. **(G)** VC-treated eyes show proliferative membranes extending over the medullary ray to the pars planar area with discontinuity(arrowheads) as a reflection of pre-existing tractional forces (scale bar: 250 μm). Magnified areas signifying area of extensive retinal fold with a pathology involving epiretinal proliferation and vitreoretinal interface disruption (scale bar: 250 μm). **(H)** Intact retinal structure with no definite epiretinal membrane or tractional forces (scale bar: 1 mm). Magnified areas signifying area of the retinal layers with no or minimal pathology relevant to the development of epiretinal proliferation or disruption of the retina (scale bar: 250 μm). **(I and J)** Immunofluorescence staining of human C-PVR cells for HLA (magenta) and nuclei (cyan) (scale bar: 50 μm). **(K)** Quantification of PVR severity scores based on clinical and OCT evaluation. Scatter plot shows individual data points with median lines. Netarsudil treatment significantly reduced PVR scores compared to VC (Mann-Whitney test; *P < 0.05).

To assess the persistence and activity of injected human C-PVR cells, immunofluorescence staining with anti-human HLA antibodies was performed. In VC eyes, HLA-positive cells were abundant and associated with sites of retinal folding, indicating active proliferation and mechanical traction. Conversely, in netarsudil-treated eyes, HLA-positive cells were sparse and largely localized to the peripapillary region or on the retinal surface, suggesting confinement to the injection site and a reduced or quiescent state of activity **(Fig. 5I and 5J, Fig. S3)**. Quantitative scoring of PVR severity based on previously published clinical and OCT criteria showed a significant reduction in the netarsudil group compared to controls **(Fig. 5K)**, supporting the therapeutic potential of netarsudil in limiting PVR progression ^15,21^.

### Netarsudil may prevent the need for re-operation in patients with PVR

In our single-surgeon series, thirty-nine surgeries performed in 29 eyes of 29 patients were included in a retrospective repeated-events analysis to evaluate for a protective effect of postoperative topical netarsudil on the need for reoperation due to PVR-related complications. Unexposed control intervals comprised 20 eyes of 20 patients who were observed without netarsudil for at least 3 months after their index operation between February 2019 and November 2024. Of these, eight eyes of eight patients (40%) underwent eleven total reoperations for PVR-C related complications during follow-up, with a median time to reoperation of 56 days. Median total follow-up time was 1 year (IQR 6m-3y). Beginning in February of 2022, patients undergoing repair of RD with PVR-C were offered treatment with topical netarsudil. Ten eyes of 10 patients were treated postoperatively with daily netarsudil 0.02% starting a median of 5 days after surgery for a median treatment duration of 109 days, and none had to have a reoperation for PVR-related complications during follow-up of at least 3 months, as a majority of redetachments occur in the first 3 months. Median total follow-up was 6 months, IQR 5 months to 1 year. The difference in follow up time between treated and untreated intervals did not reach statistical significance (p=0.09), and both groups were followed well past the highest risk period of the first 3-4 months.

Netarsudil exposure was associated with a significantly lower hazard of reoperation based on the Andersen-Gill robust score test (p = 0.01). A conservative effect size estimate was calculated using a Firth-penalized Cox model to allow a finite estimation despite observing zero reoperations in the netarsudil group. This calculation gave a hazard ratio of (HR 0.097, with a wide confidence interval of 95% CI 0.0008-0.75 estimating at least a 25% lower reoperation rate in the netarsudil-exposed intervals; p = 0.02, Firth-penalized Cox).

Mean cumulative function curves **(Fig. 6A)** illustrate that reoperations increased to approximately 0.36 per eye within the first 3 months after surgery in the control condition, further rising to 0.55 reoperations per eye by 6 months, consistent with published rates for severe PVR-C related detachments. The observed cumulative reoperation rate remained zero during netarsudil exposure; therefore, a Poisson-based upper confidence limit was estimated to illustrate the maximum cumulative reoperation risk compatible with zero observed events over a 1-year follow-up period. A swimmer plot illustrating the observation intervals and event timing of each eye over time is provided in Supplemental Figure 4.

**Figure 6.**
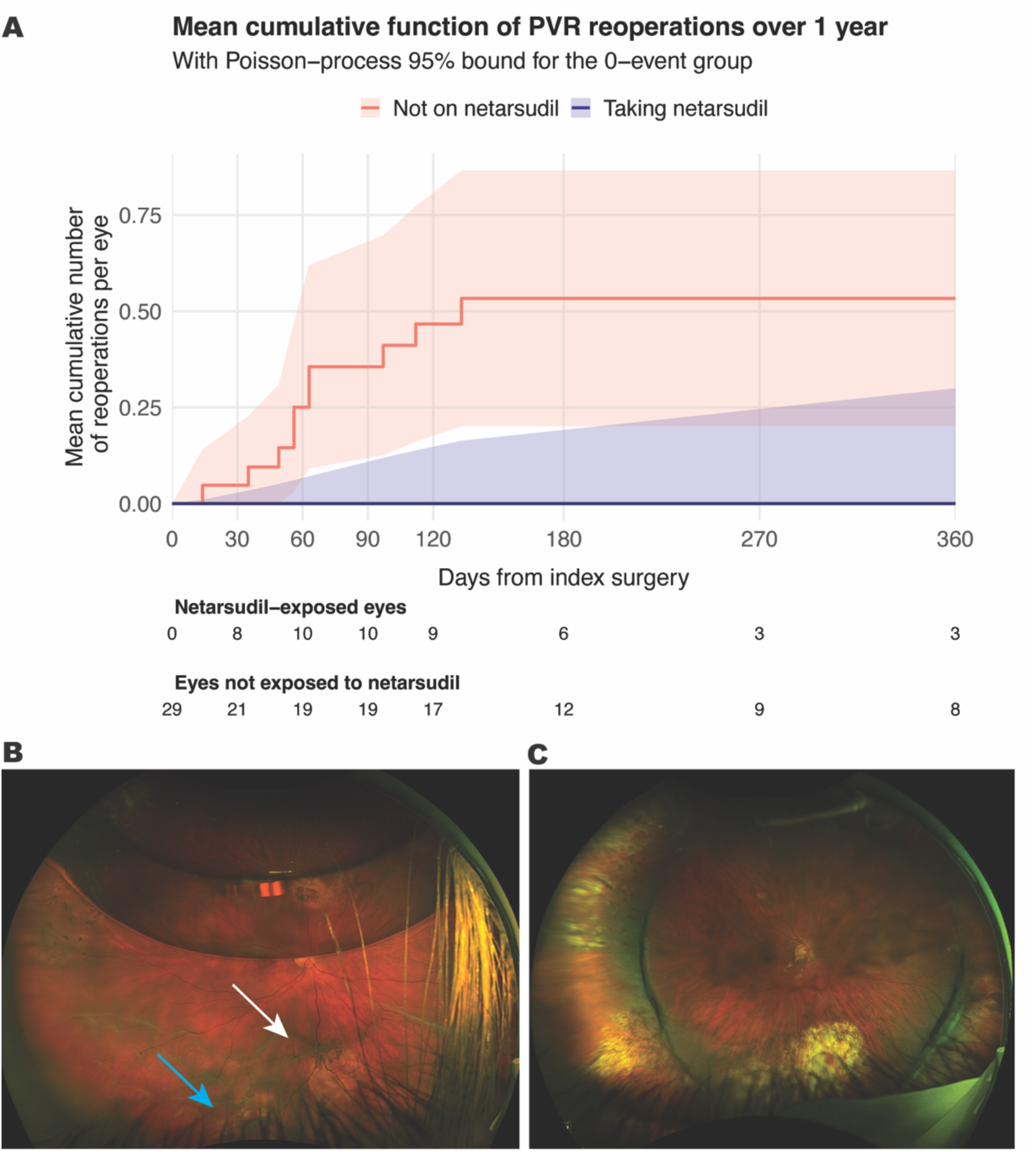
Netarsudil impact on reoperation rates, survival, and retinal structure. **(A)** Mean cumulative function plot depicting the cumulative number of reoperations per eye over time after surgery with empiric 95% CI shown for the untreated group. Approximate Poisson-based confidence limits had to be estimated for the treated group, as there were no reoperations observed. **(B)** Wide field fundus image of a recurrent RRD with PVR C (white arrow) in a patient who underwent PPV for a primary RRD. There is a retinal break inferiorly (blue arrow) and a residual C3F8 gas bubble superiorly. **(C)** Post-operative photo shows the same retina attached under silicone oil 3 months after surgery without recurrent PVR.

One patient contributed intervals to both exposure conditions due to the initiation of netarsudil only after their second retinal detachment repair in the study. This crossover is appropriately accommodated in the repeated-events analysis, which allows for time-varying exposure. For descriptive eye-level comparisons, this patient was classified in the netarsudil group, using an ever-treated categorization approach.

We tabulated a number of potential confounding factors to evaluate whether the eyes, detachments, or surgical repairs were different between the treatment and the non-treatment conditions. The median (interquartile range [IQR]) age was 68.50 (63.75, 77.25) in the group that ever received the drug and 66.00 (60.00, 72.00) (p=0.28) in the untreated patients. The youngest patient included was 43 at the time of surgery. There were no statistical differences between the groups regarding sex (p=0.52), smoking history (p=0.99), history of high myopia (p=0.42), baseline lens status (p=0.37), baseline BCVA (p=0.38), number of previous surgeries (p=0.17), RD extension (p=0.37), or PVR extension (p=0.96). All pre-operative and surgical technique data comparisons are detailed in Table 1. When analyzing all the surgeries independently, there were no statistical differences between the groups regarding the instrumentation gauge used (p=0.96), perfluoro-n-octane (PFO) use (p=0.65), the rate (p=0.95), and extent of retinectomies performed (p=0.72) or the choice of oil as tamponade (p=0.37).

**Table 1.**
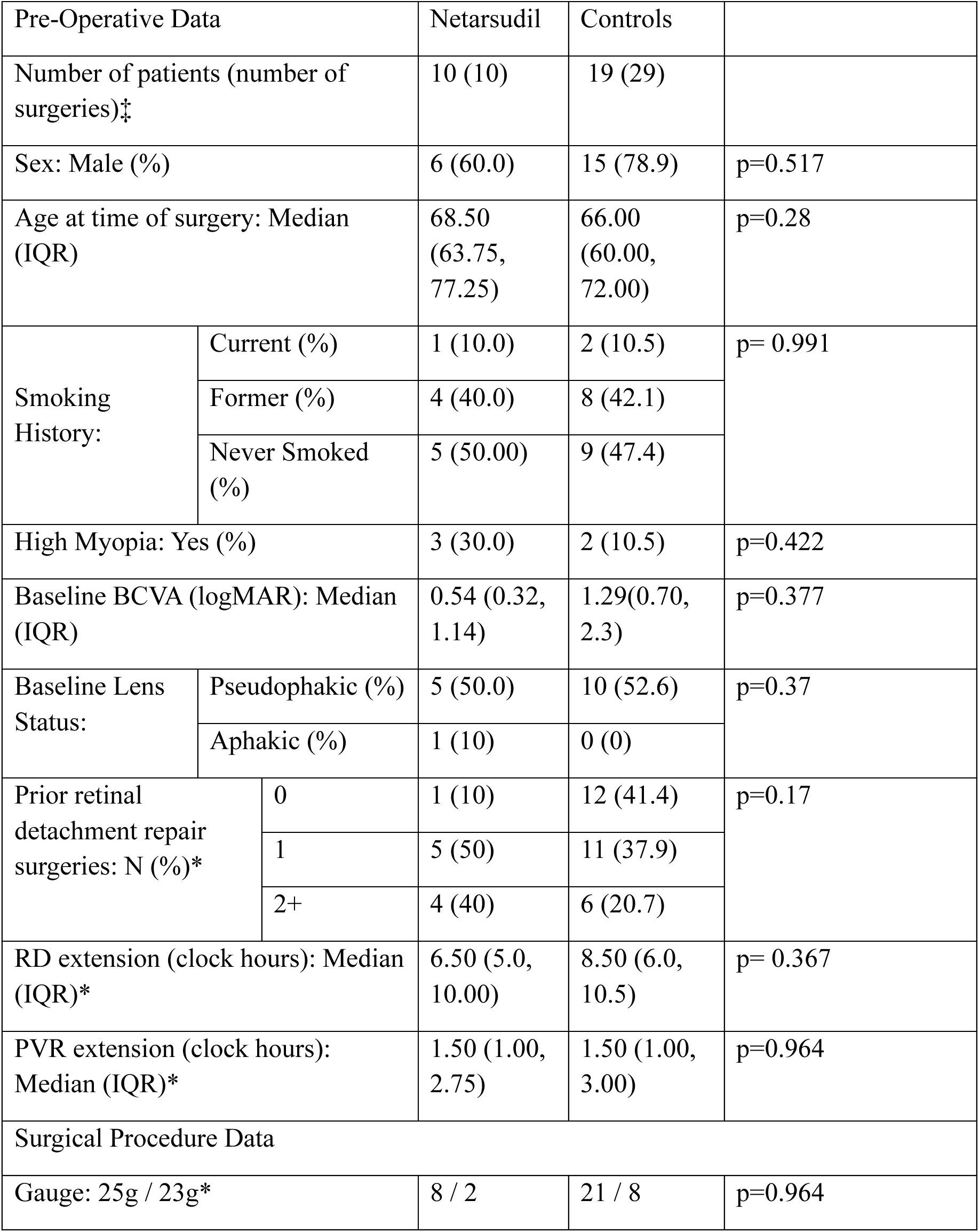
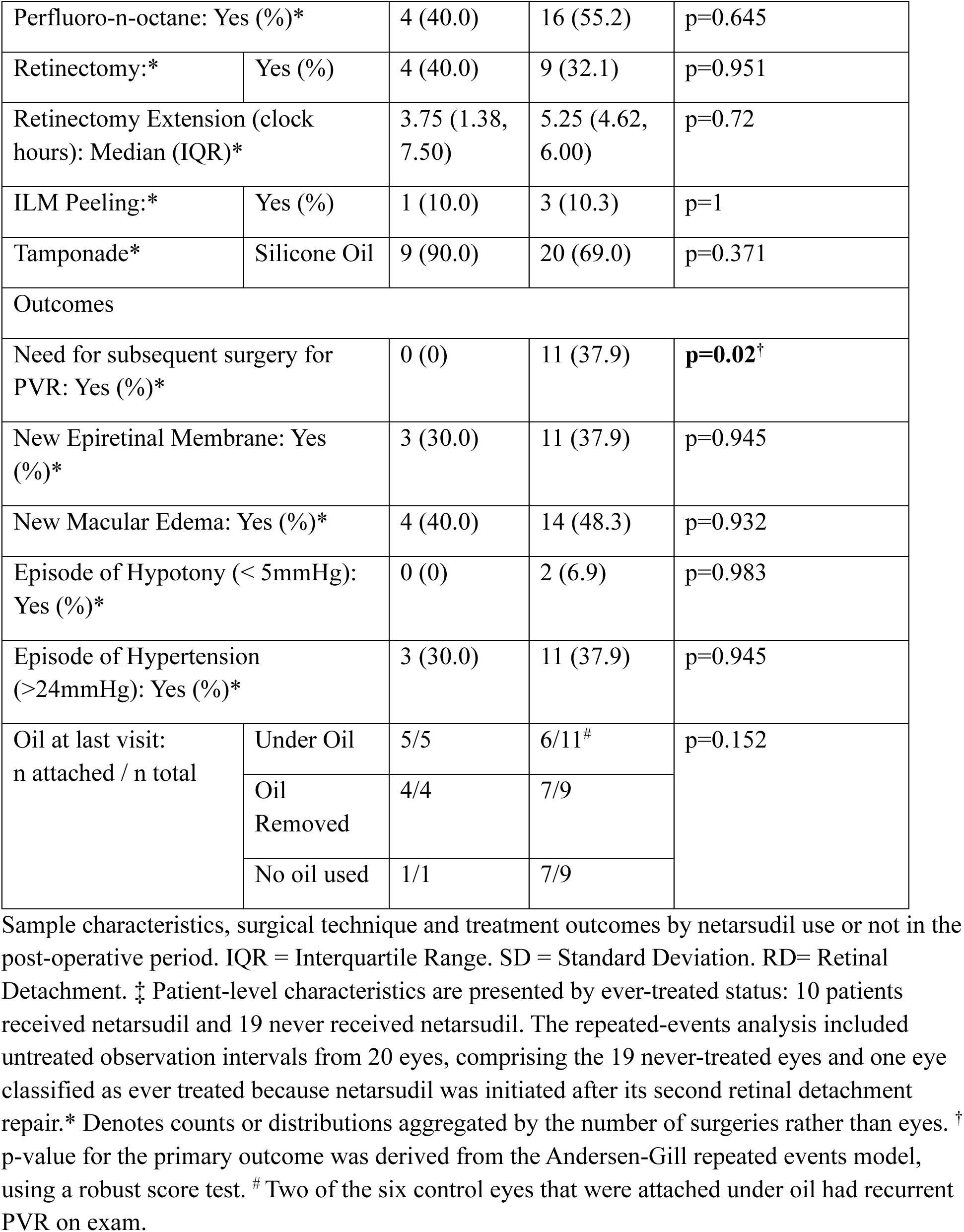
Pre-operative characteristics and surgical outcomes.

| Pre-Operative Data |  | Netarsudil | Controls |  |
|---|---|---|---|---|
| Number of patients (number of surgeries)‡ |  | 10 (10) | 19 (29) |  |
| Sex: Male (%) |  | 6 (60.0) | 15 (78.9) | p=0.517 |
| Age at time of surgery: Median (IQR) |  | 68.50 (63.75, 77.25) | 66.00 (60.00, 72.00) | p=0.28 |
| Smoking History: | Current (%) | 1 (10.0) | 2 (10.5) | p= 0.991 |
|  | Former (%) | 4 (40.0) | 8 (42.1) |  |
|  | Never Smoked (%) | 5 (50.00) | 9 (47.4) |  |
| High Myopia: Yes (%) |  | 3 (30.0) | 2 (10.5) | p=0.422 |
| Baseline BCVA (logMAR): Median (IQR) |  | 0.54 (0.32, 1.14) | 1.29(0.70, 2.3) | p=0.377 |
| Baseline Lens Status: | Pseudophakic (%) | 5 (50.0) | 10 (52.6) | p=0.37 |
|  | Aphakic (%) | 1 (10) | 0 (0) |  |
| Prior retinal detachment repair surgeries: N (%)* | 0 | 1 (10) | 12 (41.4) | p=0.17 |
|  | 1 | 5 (50) | 11 (37.9) |  |
|  | 2+ | 4 (40) | 6 (20.7) |  |
| RD extension (clock hours): Median (IQR)* |  | 6.50 (5.0, 10.00) | 8.50 (6.0, 10.5) | p= 0.367 |
| PVR extension (clock hours): Median (IQR)* |  | 1.50 (1.00, 2.75) | 1.50 (1.00, 3.00) | p=0.964 |
| Surgical Procedure Data |  |  |  |  |
| Gauge: 25g / 23g* |  | 8 / 2 | 21 / 8 | p=0.964 |
| Perfluoro-n-octane: Yes (%)* |  | 4 (40.0) | 16 (55.2) | p=0.645 |
| Retinectomy:* | Yes (%) | 4 (40.0) | 9 (32.1) | p=0.951 |
| Retinectomy Extension (clock hours): Median (IQR)* |  | 3.75 (1.38, 7.50) | 5.25 (4.62, 6.00) | p=0.72 |
| ILM Peeling:* | Yes (%) | 1 (10.0) | 3 (10.3) | p=1 |
| Tamponade* | Silicone Oil | 9 (90.0) | 20 (69.0) | p=0.371 |
| Outcomes |  |  |  |  |
| Need for subsequent surgery for PVR: Yes (%)* |  | 0 (0) | 11 (37.9) | **p=0.02**<sup>†</sup> |
| New Epiretinal Membrane: Yes (%)* |  | 3 (30.0) | 11 (37.9) | p=0.945 |
| New Macular Edema: Yes (%)* |  | 4 (40.0) | 14 (48.3) | p=0.932 |
| Episode of Hypotony (< 5mmHg): Yes (%)* |  | 0 (0) | 2 (6.9) | p=0.983 |
| Episode of Hypertension (>24mmHg): Yes (%)* |  | 3 (30.0) | 11 (37.9) | p=0.945 |
| Oil at last visit:<br>n attached / n total | Under Oil | 5/5 | 6/11<sup>#</sup> | p=0.152 |
|  | Oil Removed | 4/4 | 7/9 |  |
|  | No oil used | 1/1 | 7/9 |  |

IQR = Interquartile Range. SD = Standard Deviation. RD= Retinal Detachment. ‡ Patient-level characteristics are presented by ever-treated status: 10 patients received netarsudil and 19 never received netarsudil. The repeated-events analysis included untreated observation intervals from 20 eyes, comprising the 19 never-treated eyes and one eye classified as ever treated because netarsudil was initiated after its second retinal detachment repair.\* Denotes counts or distributions aggregated by the number of surgeries rather than eyes. <sup>†</sup> p-value for the primary outcome was derived from the Andersen-Gill repeated events model, using a robust score test. <sup>#</sup> Two of the six control eyes that were attached under oil had recurrent PVR on exam.

Among the reoperations observed during follow up, all of which occurred in untreated eyes, 9 were necessary due to a PVR-induced recurrent RRD, while 2 had PVR-induced hypotony without RRD. C3F8 was chosen as a tamponade in 1 of the surgeries in the netarsudil group and in 9 of the control surgeries, while silicone oil was chosen in 9 of the netarsudil group and 20 of the controls (p=0.37). Four (40%) surgeries in the netarsudil group and 9 (45%) in the control group had the oil removed before the final follow-up.

While the study surgeon completed a vitreoretinal surgery fellowship in 2010 and was not aware of any changes in PVR repair techniques after February 2022 (when netarsudil was first offered) compared to the period between 2019-2022, it is conceivable that subconscious alterations in techniques accounted for the zero re-operation rate in the netarsudil group, which were all operated on after 2022. We found that the number of re-operations in patients with PVR C not receiving netarsudil between February 2022 and November 2024 was three of 11 eyes in 11 subjects (27%; p-value=0.68 compared to the 40% rate of re-operations for PVR in the control group between February 2019-2022). Notably, these 11 patients who did not take the netarsudil that was offered after February 2022 cited various reasons such as cost and off-label use.

Among the subjects that received netarsudil, five started the drug one day after the surgery, while one started 38 days after the procedure due to a corneal epithelial defect in a penetrating keratoplasty corneal graft; the other four patients started 9 or 10 days after the vitrectomy (Median = 5 days / IQR =1.0,9.8). The median (IQR) time of use was 109.00 (104.00, 126.00) days after the surgery, and no patient had to stop the drug due to adverse effects. During the post-operative period, no patient in the netarsudil group developed hypotony (<5 mmHg), compared to two episodes in the control group (p=0.98). Both instances of hypotony occurred in consecutive surgeries of the same patient and were related to extensive PVR development in the retinal periphery, with presumed tractional effect on the ciliary body. There were three episodes of ocular hypertension (>24 mmHg) reported in patients who received netarsudil, and 11 in the control group (p=0.95). None of the episodes in the netarsudil group exceeded 30mmHg and all were controlled within a week with drops. One of the patients in the netarsudil group had a previous history of ocular hypertension and developed an episode of hypertension three months after the surgery. All other episodes of ocular hypertension in the netarsudil group happened within one week after the surgery and occurred in patients who had not started the drug up to that point. On the other hand, six of the patients in the control group had IOP recorded over 30mmHg. The maximum IOP recorded was 52 mmHg. All the ocular hypertensive episodes in the control group were successfully treated with hypotensive drops within one week. Three episodes of new epiretinal membrane were observed in the netarsudil group compared with 11 in the control group (p=0.95). There were no episodes of endophthalmitis in either of the groups. Best corrected visual acuity between the two groups did not differ at the final post-operative follow-up (p=0.46).

## DISCUSSION

PVR remains a major cause of failure after retinal detachment repair, with limited pharmacologic options ^1,2,5^. This study brings together patient-derived cellular and tissue models, single-cell profiling, an experimental rabbit model, and exploratory clinical observations to investigate ROCK inhibition as a therapeutic strategy. Netarsudil-associated reductions in C-PVR cell proliferation, migration, and contraction, together with reduced explant outgrowth and experimental disease severity, provide complementary preclinical evidence of antifibrotic activity. The exploratory association between topical treatment and fewer PVR-related reoperations warrants prospective investigation, but differences in drug delivery and the clinical study design limit direct extrapolation between experimental and patient outcomes.

RhoA/ROCK signaling regulates cytoskeletal organization, migration, and contraction and participates in TGF-β-associated fibrotic responses ^8,19^. ROCK-mediated regulation of actomyosin dynamics provides a mechanistic rationale for effects on cellular migration and contractility, although downstream pathway engagement was not established across all experimental systems. The functional assays suggest that ROCK inhibition affects several behaviors contributing to PVR progression. Previous work showed that methotrexate reduced C-PVR cell proliferation and induced apoptosis, with no detectable effect on migration ^22^. The additional effects of ROCK inhibitors on migration and contraction therefore provide a complementary therapeutic rationale.-Previous work implicating ROCK in focal adhesion dynamics and demonstrating reduced extracellular matrix expression in RPE models, including TGF-β2-induced type I collagen, further supports the relevance of this pathway to fibrotic membrane biology ^23–25^.

Single-cell profiling provides transcriptional context for the functional findings, although the patient-membrane and cultured-cell datasets address distinct biological questions. Profiles from patient membranes characterize cellular heterogeneity in diseased tissue, whereas treatment-associated differences in cultured C-PVR cells identify candidate molecular programs responsive to netarsudil. Differences in matrix-associated expression programs and their relative representation are compatible with an antifibrotic response but do not establish conversion between cellular states. Because these analyses represent defined sampling points, changes in state proportions could also reflect differential survival or recovery of subpopulations. Longitudinal profiling and orthogonal validation will be needed to link these transcriptional associations to specific functional mechanisms.

Patient-membrane explants and the rabbit model extend these observations across experimental systems. Reduced explant outgrowth supports an effect in patient-derived tissue, where the measured response integrates proliferation, migration, and survival. In the established rabbit model ^15,21^, intravitreal netarsudil attenuated PVR severity and membrane formation, with imaging and histology supporting preservation of retinal architecture. HLA staining provided complementary information about the distribution of transplanted human cells. Together, these findings support antifibrotic activity under direct experimental exposure. However, explant outgrowth is a composite endpoint, and the induced rabbit model incompletely reproduces the inflammatory, surgical, and retinal-break heterogeneity of human PVR.

Netarsudil’s existing ophthalmic use makes it a practical candidate for further investigation. ROCK inhibition is an established therapeutic approach in glaucoma ^26,27^, and topical administration could reduce the procedural burden of adjunctive treatment if efficacy in PVR is confirmed. The principal translational gap is drug exposure: cellular, explant, and rabbit experiments involved direct administration, whereas patients received topical drops. Posterior-segment concentrations of netarsudil and its pharmacologically active metabolites after topical dosing were not established in this study. Pharmacokinetic and pharmacodynamic studies in postoperative eyes are therefore needed to determine whether topical dosing can achieve exposures associated with the experimental antifibrotic response and to define an appropriate dosing and safety window.

The exploratory clinical cohort provides an initial signal in patients undergoing surgery for established grade C PVR, a population at substantial risk of recurrent complications ^28–30^. No PVR-related reoperations were observed among treated patients, compared with a 40% reoperation proportion in controls. Although this observation supports prospective evaluation, the nonrandomized comparison and absence of treated events prevent a precise estimate of the magnitude or causal nature of any benefit. Reoperation is a clinically important endpoint but should not be equated with the absence of recurrent PVR. The clinical question concerns recurrent PVR-related surgical failure after repair; prevention in eyes without established PVR represents a separate application. Topical treatment may offer an administration advantage over repeated intravitreal regimens, but comparative efficacy requires direct evaluation.

Several mechanistic questions remain. Although the functional assays demonstrate treatment-associated reductions in PVR-relevant behaviors, the sequence connecting ROCK pathway inhibition to these phenotypes was not comprehensively established across experimental systems. Time-resolved measurements of actomyosin-related pathway activity, together with orthogonal protein-level validation of selected single-cell findings, would clarify the relationships among target engagement, transcriptional responses, and function. Concentration-response and viability assessments would also help distinguish selective antifibrotic activity from nonspecific reductions in cellular fitness.

The clinical observations are limited by the nonrandomized study design and small cohort size. Differences in baseline disease severity, surgical management, concomitant therapy, and duration or completeness of follow-up could confound the association between treatment and reoperation. Variation in the timing of postoperative netarsudil initiation requires particular attention, because exposure classification and follow-up must account for the period between surgery and treatment initiation to minimize time-related bias. PVR-related reoperation is clinically meaningful but may not capture all recurrent membrane formation or disease progression, and decisions to reoperate can vary. With no reoperations among treated eyes, the estimated treatment association remains imprecise. These findings should therefore be considered hypothesis-generating, not evidence of established clinical efficacy.

Overall, the preclinical findings support further investigation of ROCK inhibition for PVR, and the clinical observations provide an exploratory rationale for evaluating topical netarsudil as an adjunct to surgery. Progress toward clinical application requires demonstration of relevant posterior-segment exposure, definition of dosing and safety, and prospective controlled assessment of PVR-related reoperations. A staged translational approach, including ocular exposure and target-engagement studies followed by prospective controlled evaluation, is needed before clinical efficacy can be established.

## MATERIALS AND METHODS

### Patient-derived primary cell culture

In vitro assays were conducted using a primary cell culture derived from a patient with PVR that had been previously established and referred to as C-PVR cells ^22^. The culture medium of C-PVR cells (C-PVR medium) is composed of MCDB 131 medium (Gibco^TM^, 10372019), which was supplemented with EGM-2 Endothelial Cell Growth Medium-2 SingleQuots Supplements and Growth Factors (Lonza, CC-4176), 1% L-glutamine, 100 U/mL (Gibco^TM^, 35050-061), Penicillin 10,000 Units/mL, Streptomycin 10,000 μg/mL (Gibco^TM^, 15140-122), and 10% fetal bovine serum (R&D Systems, S11150). C-PVR cells were maintained in an incubator under standard human cell conditions (i.e., 37°C, 5% CO2).

### Cytotoxicity and cell proliferation assays

A 96-well plate was seeded with 10,000 C-PVR cells per well in C-PVR medium. Subsequently, cells were exposed to different concentrations (0.1 to 10 μM) of netarsudil (Abcam, Inc., ab287030). C-PVR cells not exposed to any treatment (NTC) and cells exposed to vehicle (VC; DMSO; MilliporeSigma, D2653-5X10ML) as controls. The plates were incubated at 37°C in a 5% CO₂ atmosphere for 24, 48, and 72 hours. To assess lactate dehydrogenase (LDH) release, 100 μL of supernatant was taken from each well and analyzed using Cytotoxicity Detection Kit (LDH; Roche,11644793001) in accordance with the manufacturer’s instructions. The LDH release percentage was calculated for each incubation time relative to the NTC, representing the maximum LDH release. The levels of LDH were quantified by measuring the absorbance at a wavelength of 450 nm. To assess the inhibitory potential of netarsudil on C-PVR cells, the fluorescence-based CyQUANT Cell Proliferation Assay Kit (Invitrogen, C7026) was employed. The percentage of proliferation was quantified relative to the NTC for each time point at concentrations that did not exhibit cytotoxic effects in the LDH assay. Absorbance and fluorescence (ex/em: 563/587 nm) readings were conducted on a Synergy H1 Hybrid Microplate Reader (BioTek).

### Apoptosis and necrosis assay

To assess apoptosis and necrosis, we used the RealTime-Glo™ Annexin V Apoptosis and Necrosis Assay (Promega, JA1011) according to the manufacturer’s instructions. Briefly, cells were seeded in 96-well plates at a density of 10,000 cells per well and allowed to adhere overnight. The following day, cells were treated with netarsudil at 2.5, 5.0 and 10.0 μM, VC, and NTC for 2, 4, 10, 24, 40, 48, and 72 hours. Assay reagents, consisting of the RealTime-Glo™ Annexin V reagent and necrosis detection reagent, were added directly to the culture medium at the recommended concentrations. Apoptosis was measured by luminescence, reflecting Annexin V binding to externalized phosphatidylserine, while necrosis was assessed by fluorescence, indicating loss of membrane integrity. Luminescence and fluorescence signals were recorded at the indicated time points using a Synergy H1 Hybrid Microplate Reader (BioTek). Background signals were subtracted using blank control wells. Data were normalized to baseline readings and expressed as relative luminescence/fluorescence units (RLU/RFU).

### Wound healing assay

Wound healing assays were conducted in 24-well plates. A total of 150,000 cells were seeded per well, and immediately exposed to 2.5 µM netarsudil, vehicle (VC, DMSO), or not exposed to any treatment (NTC). After 24 hours, a cell-free gap was created using a 200 µL tip, which was used to scratch the cell monolayers in each well. At this point, the treatments were replaced, and the wound size was photographically monitored. Images were acquired using an EVOS M7000 Imaging System (Invitrogen) at 0-, 3-, 6-, 9-, 12-, 15-, and 18-hours post-scratch. For quantification, the wound area was measured using the Wound Healing Size Tool plugin in Fiji (ImageJ) ^31^. Wound closure was normalized to the initial wound area at 0 hours, and data were expressed as the percentage of wound closure.

### Collagen-based contraction assay

The effect of netarsudil on C-PVR cell contraction was evaluated using a CytoSelect™ 48-Well Cell Contraction Assay Kit (Floating Matrix Model; Cell Biolabs, CBA-5021) following the manufacturer’s instructions. A total of 50,000 cells were seeded per well in a 48-well plate. Cell-free wells were used as a control to represent the maximum surface area of the gel. Following the polymerization of the gel, 500 μL of medium was added containing the following treatment: (i) netarsudil, (ii) TNF-α, (iii) TGF-β2, (iv) TGF-β2+netarsuil. Non-treated cells (NTC) and vehicle-treated cells (VC, DMSO) were used as controls. The treatments were partially refreshed (250 µL) every 24 hours. After 72 hours, the medium was completely removed, and photographs of each well were captured using an M60 stereo microscope equipped with an IC90 E (Leica Microsystems). The surface area of each gel was quantified using the ImageJ software. The data were expressed as relative percentages of gel surface area (normalized against the cell-free well control).

### Patient-derived explants

Membranes obtained from the patient following recent surgical procedures were dissected into small fragments and embedded in Cultrex (basement membrane matrix, R&D Systems, BME001-05). C-PVR medium containing 12% bovine calf serum was added and incubated at 37°C with 5% CO₂. Media was replaced every 48 hours. Treatments were performed after ensuring the growth of the explants. Each explant was photographed using an inverted cell culture phase contrast microscope CKX53 equipped with an EP50 digital camera (Olympus) to establish a baseline for the observation period. Images were taken every two or three days, up to twelve days. Explant growth was quantified using Fiji (ImageJ2 v2.16.0/1.54g ^32^). Images were first converted to an 8-bit grayscale, and the scale was set according to the image capture source. Background subtraction was performed by selecting a representative background region, measuring its mean gray value, and subtracting this value from the image. To enhance boundary detection, the “Find Edges” function was applied. Growth area was then segmented using the thresholding tool, with default values applied unless manual adjustment was needed to reduce background noise. Finally, the explant growth area was measured based on the thresholded region, providing a quantification of the expanding tissue. To standardize growth measurements across tissue explant samples, we normalized the area at each time point relative to the final measured area. The area at each time point was expressed as a percentage of the final area, which was assumed to represent 100% growth. This approach allowed for a direct comparison of growth dynamics across different explants, independent of initial size variations.

### PVR Induction and Netarsudil Administration in Rabbits

Proliferative vitreoretinopathy (PVR) was induced in male New Zealand White rabbits (2.3 kg, 6-8 weeks old; Charles River Laboratories, Wilmington, MA) using a previously established method ^15,21^. In brief, a 0.15 mL intravitreal injection of perfluoropropane gas (C₃F₈; Alcon, 8065797102) was administered 3 mm posterior to the limbus to displace the vitreous. After three days, intraocular pressure was reduced via partial gas removal. Subsequently, a mixture of 1 × 10⁶ C-PVR cells in 0.1 mL of balanced salt solution (BSS) was injected intravitreally, followed by 50 µL of the netarsudil formulation (75 µM in BSS, yielding ∼2.5 µM vitreous concentration). A second dose of the netarsudil formulation (50 µL) was administered one week later. Disease severity was evaluated two weeks after C-PVR cell injection using a composite PVR grading scale based on indirect ophthalmoscopy, OCT, and fundus imaging. Scoring was performed independently by two masked ophthalmologists using the Fastenberg scale **(Table S2)** and a PVR grading system, previously developed by our group ^21^. Imaging and histological assessments were conducted using established protocols.

### Histological Analyses of Rabbit Eyes

Histological analysis of rabbit eyes was performed as previously described ^15,21^. Following euthanasia, enucleated eyes were fixed in Davidson’s fixative (Millipore Sigma, H0290) at room temperature for 24 hours. A 1 × 1 mm scleral incision was made to enhance fixative penetration, and eyes were immersed in fixative for an additional 24 hours. Tissues were then transferred to 70% ethanol for at least 24 hours before paraffin embedding. Serial sections (5 μm) were cut and stained with hematoxylin and eosin (H&E) for morphological evaluation of the retina and associated PVR membranes.

To confirm the presence of human cells within the rabbit vitreous cavity, a specific antibody against human antigen was used: rat anti-human leukocyte antigen (HLA) Class I Xenograft marker antibody (1:200; SM2012P Acris, OriGene Technologies, Inc., Rockville, MD). Slides were washed with PBS and permeabilized with 0.5% Triton X-100 in PBS for 5 min and blocked (10% goat serum in PBS) for 1 hour at RT. The primary antibody was prepared in antibody dilution buffer (5% goat serum) and incubated overnight at 4 °C. Slides were washed with PBS and incubated with goat anti-rat and goat anti-mouse Alexa Fluor 594 secondary antibody, respectively (1:300; Life Technologies, A-11007, A-11005) for 2 hours at RT. Slides were mounted and visualized using Prolong Gold Antifade Reagent with DAPI (Invitrogen, P36935). Images were obtained using a Leica SP8 Confocal Laser Microscope (Weitzlar, Germany).

### Immunoblotting

Protein levels were determined using standard Western blotting practices. Briefly, cell lysates were prepared, and the protein concentration was measured using a BCA assay. Equal amounts of protein were separated using SDS-PAGE and transferred to PVDF membranes. Membranes were blocked with 5% non-fat dry milk for 1 hour at room temperature and then incubated with primary antibodies **(Table S3)** overnight at 4°C. After washing, membranes were incubated with HRP-conjugated secondary antibodies (Table S3) for 1 hour at room temperature. Detection was performed using an enhanced chemiluminescence mixture Clarity^TM^ Western ECL Substrate (Bio-Rad, 1705061), and the blots were imaged using a digital imaging system iBright FL1500 (Thermo Fisher Scientific;). Bands were quantified using ImageJ software.

### Single-Cell RNA Sequencing (scRNA-seq) Data Acquisition and Processing

Published scRNA-seq data from freshly obtained proliferative vitreoretinopathy (PVR) membranes were downloaded as a Seurat object from the Single Cell Portal (SCP2582)^15^ and reanalyzed to characterize ROCK pathway gene expression. ScType^16,33^ was applied to scaled expression data, with marker scores aggregated by cluster. Related immune labels were consolidated after manual review. UMAP and ridge plots were generated in Seurat (v5.2.1)^34^.

Cultured PVR (C-PVR) cells were exposed to vehicle or 2.5 μM netarsudil for 48 hours and dissociated with TrypLE (Gibco). Two biological replicates per condition were processed independently. The Boston University Single-Cell Sequencing Core prepared 10x Genomics Chromium GEM-X 3′ libraries and sequenced them on an Illumina NextSeq 2000. Cell Ranger was used for demultiplexing, alignment, cell calling (expected cell count, 10,000), and unique molecular identifier (UMI) quantification. Droplets with ≥500 UMIs underwent SCTK-QC screening for low-quality cells, doublets, and ambient RNA^35^.

Replicate count matrices were merged in Seurat (v5.2.1) in R (v4.4)^34^. Data were normalized with SCTransform and visualized by UMAP after principal component analysis. ScType^16,33^ scored clusters against combined eye and immune marker sets. Clusters were initially assigned their highest-scoring label or designated Unknown if that score was below one-quarter of the number of cells in the cluster. The scType-assigned labels were manually reviewed and revised to reflect the closest transcriptional state supported by the observed gene expression profiles, without implying definitive cell identity or origin.

Cell-state fractions were calculated within each condition and compared using Fisher’s exact tests with Benjamini–Hochberg false discovery rate correction (https://www.jstor.org/stable/2346101). Seurat FindMarkers was used to compare gene expression between conditions at log fold-change thresholds of 0, 0.5, and 1 (adjusted P < 0.05). Genes were mapped to Entrez IDs with biomaRt (v2.60) for Gene Ontology enrichment with clusterProfiler (v4.12.6). MSigDB Hallmark gene sets were tested on log fold-change-ranked genes using fgsea (v1.30)^36,37^ (adjusted P < 0.05). CellChat (v1.6.1)^38^ inferred condition-specific ligand–receptor networks for secreted signals in CellChatDB.human, with a minimum of 10 cells per group. The Supplementary Methods file provides the full scRNA-seq methodology and lists the original scType annotations alongside the revised labels.

### Clinical study design and participants

Patients given netarsudil (and controls) were cared for by a single surgeon (EHS) at University of Iowa Healthcare, which granted Institutional Review Board approval (#202404354) for this retrospective, interventional, comparative analysis. All patients underwent standard of care surgery for PVR removal between February 2019 and November 2024, and were included if they had primary rhegmatogenous retinal detachment (RRD) repairs associated with proliferative vitreoretinopathy (PVR) grade C, re-detachment due to PVR C, and/or PVR C removal due to secondary hypotony (e.g. epiciliary membrane removal) in the post operative period of an RRD repair. The study evaluated the time-dependent rate of re-operation for PVR-related complications after surgery for RRD with PVR C. The index time of observation for at-risk eyes was the first eligible surgery performed by the study surgeon EHS. Events were defined as a reoperation for a complication of recurrent PVR, such as redetachment. Each eye contributed multiple observation intervals spanning from the index surgery through subsequent eligible procedures or censoring at the last follow-up. Patients were found using the RedCap and SlicerDicer systems at the University of Iowa in addition to the surgeon’s personal surgical logbook.

The study group used one drop of netarsudil 0.02% every evening in the affected eye for at least three months after their surgery, and the risks, benefits, and alternatives of this including the off-label nature were explained in detail to patients. Patients paid out of pocket for the medication if their insurance did not cover it. The control group had standard of care surgery and post-operative care without the addition of netarsudil. Netarsudil was offered to all patients with PVR grade C after February 2022. There were no other differences between these two groups, as the surgeon maintained the same standard of care with or without netarsudil.

Exclusion criteria consisted of patients that had PVR grade A or B; did not receive pars plana vitrectomy (PPV); had surgery performed before February 2019; had active uveitis; had less than 3 months post-operative follow-up; had a history of ocular trauma, previous use of rho kinase inhibitors, presence of tractional detachment, proliferative or severe non proliferative diabetic retinopathy, retinal neovascularization, sub-retinal neovascular membranes, ocular tumors, radiation retinopathy, inherited vitreoretinal dystrophies, retinal detachment associated with macular holes; and those who received intravitreal methotrexate. If both eyes were eligible, only the eye that received surgery first was included.

### Surgical Technique

All patients underwent 23 or 25-gauge three port vitrectomy performed by the same experienced surgeon (EHS). A complete vitrectomy with posterior hyaloid detachment and vitreous base shaving was performed in all patients. Mechanical removal of all PVR membranes was performed. Perfluoro-n-octane was used to open and stabilize the retina when the membrane removal was not possible under balanced saline solution. If sub-retinal bands were observed, a retinotomy was performed if the membrane could not be accessed through the retinal tear. Relaxing retinectomy was performed when the retina would not flatten despite removal of PVR membranes. Subretinal fluid was preferentially drained through a retinal tear or through the retinectomy if present. When drainage was not possible through the tear and no retinectomy was performed, a posterior small retinotomy was created. Scleral buckle (encircling 42 band with 70 Watzke sleeve) was placed to support the vitreous base in patients with recurrent retinal detachment from PVR grade C and/or when an inferior relaxing retinectomy was performed. Epiretinal membrane (ERM) was peeled if present. Three rows of adjoining laser spots were applied in the margin of each identified retinal tear, retinotomy and the retinectomies. Silicone oil was used in patients who had a questionable ability to position prone post-op and for those with inferior relaxing retinectomies.

### Statistical Analysis

For in vitro and ex vivo data, Western blot analysis was quantified using ImageJ software V13 (National Institutes of Health). Data are presented as the mean ± standard deviation. Statistical analysis was conducted using GraphPad Prism version 10.1.2, considering p-values below 0.05 as statistically significant. Initially, residuals of raw data were assessed for normality to determine whether parametric or nonparametric tests were appropriate. Parametric data were analyzed using the Student’s t-test or one-way/two-way ANOVA, followed by Tukey’s test for multiple comparisons unless specified otherwise. Pearson’s rank correlation coefficient was used to assess rank correlation. For nonparametric data, the Mann-Whitney U test, Kruskal-Wallis test, or Friedman’s test was applied, with Dunn’s post-hoc test used for multiple comparisons.

Statistical analysis for clinical data was performed using R (www.r-project.org, version 4.2.1) with the *survival* and *coxpf* libraries. For the primary clinical analysis, reoperations were analyzed as recurrent events using an Andersen-Gill model with total time since index as the timescale. Risk intervals were constructed using counting process form, and netarsudil exposure was modeled as a time-varying covariate that was persistent after initiation. Robust sandwich variance estimates clustered by eye were used to account for within-eye correlation. Because no events occurred during netarsudil exposure, the standard partial likelihood exhibited separation. Therefore, inference was based on the robust score test, supplemented by nonparametric mean cumulative function curves, and a Firth-penalized Cox model was fit as a sensitivity analysis to obtain finite hazard ratio estimates. Given the limited sample size, models were restricted to treatment as the primary covariate. Treatment and control groups were evaluated for similarity using the Fischer Exact test for categorical variables, Student’s t-test for normally distributed continuous data, or a Wilcoxon rank sum test for non-normally distributed continuous data.

### Study approval

All procedures involving secondary use of human tissues and clinical specimens were approved by Mass General Brigham (protocol no. 2022P002602). Written informed consent was obtained from all participants or legal guardians. Animal experiments using rabbits were approved by the MEE Institutional Animal Care and Use Committee (IACUC) (protocol no. 2021N000101). The clinical pilot study was approved by the University of Iowa Healthcare Institutional Review Board (protocol no. 202404354) and conducted following Good Clinical Practice guidelines.

Written informed consent was obtained from all participants.

## Acknowledgments

We would like to thank the Enterprise Research Infrastructure & Services (ERIS) at Mass General Brigham for their in-depth support and for the provision of computational resources.

## Funding

This research was supported by the Department of Defense US Army Medical Research Acquisition Activity: HT9425-23-1-0527 (LAK, MOH).

This work was supported in part by NEI Grant R01-EY035435 (EHS) for studies on retinal fibrosis, and by VA Career Development Award IK1RX005029 (EFL) that supported statistical analysis and clinical modeling.

This work is also supported, in part, by Michel Plantevin and the Monte J. Wallace Ophthalmology Chair in Retina (LAK).

## Author contributions

Conceptualization: LAK, MOH, EHS

Methodology: JS-S, BB, WPM, AU, RL, AV, KWB, EHS, MOH, LAK

Investigation: JS-S, BB, WPM, AU, RL, DA, DYS, AV, KWB, YA-R, MS, BAK, ALG, KB

Visualization: JS-S, BB, WPM, AU, RL, DA, DYS, AV, KWB, YA-R, MS, BAK, ALG, NR, EJR, DE, EFL, EHS, MOH, LAK

Funding acquisition: EFL, MOH, LAK, EHS Project administration: MOH, LAK

Supervision: MOH, LAK, EHS

Writing – original draft: JS-S, MOH, BB, NR, WPM, EHS, LAK

Writing – review & editing: JS-S, BB, WPM, AU, RL, DA, DYS, AV, KWB, YA-R, MS, BAK, ALG, JFR, RPB, KB, PP, AL, NR, EJR, DE, EFL, EHS, MOH, LAK

