## Supplemental Table S2 for "Targeting Rho Kinase in Proliferative Vitreoretinopathy with Netarsudil: Multimodal Preclinical Evidence and Exploratory Clinical Application"

**Table S2**. Predictors of Subsequent Surgery Due to PVR.

| **Patient Data** | | **No Reoperation** | **Reoperated** | |
|---|---|---|---|---|
| Total | | 22 | 8 | |
| Netarsudil Used in the Post-Operative Period (%) | | 10 (45.45) | 0 (0) | p=0.029 |
| Sex: Male (%) | | 15 (68.2) | 4 (50.0) | p=0.39 |
| Age at time of surgery: Median (IQR) | | 67.00 (63.00, 72.00) | 68.0 (67.00, 72.00) | p=0.672 |
| Smoking History: | Current (%) | 2 (9.1) | 0 (0) | p=0.352 |
| | Former (%) | 10 (45.5) | 6 (75.0) | |
| | Never Smoked (%) | 10 (45.5) | 2 (25.0) | |
| High Myopia: Yes (%) | | 5 (22.7) | 1 (12.5) | p=1 |
| Number of Previous Surgeries for PVR: Median (IQR) | | 1 (0.25,2.00) | 1 (0.00,1.00) | p=0.342 |
| Pre-Operative BCVA (logMAR): Median (IQR) | | 0.79 (0.54,2.12) | 2.3 (1.60,3.30) | p=0.277 |
| Pre op Lens Status: | Aphakic (%) | 3 (13.6) | 1 (12.5) | p=1 |
| | Pseudophakic (%) | 11 (50.0) | 4 (50.0) | |
| | Phakic (%) | 8 (36.4) | 3 (37.5) | |
| RD extension (clock hours): Median (IQR) | | 7.00 (5.12,9.50) | 9.00 (6.62,11.38) | p=0.336 |
| PVR extension (clock hours): Median (IQR) | | 1.50 (1.00,2.50) | 3.00 (2.50,3.00) | p=0.072 |
| **Surgical Procedure** | | | | |
| Gauge: 25g / 23g | | 15 / 7 | 5 / 3 | p=1 |
| Perfluoro-n-octane: Yes (%) | | 13 (59.1) | 3 (37.5) | p=0.417 |
| Retinectomy: | Yes (%) | 9 (40.9) | 3 (37.5) | p=1 |
| | Previous (%) | 1 (4.5) | 0 (0) | |
| | No (%) | 12 (54.5) | 5 (62.5) | |
| Retinectomy Extension (clock hours): Median (IQR) | | 5.5 (4.12,6.00) | 5.00 (3.25,5.25) | p=0.492 |
| Concurrent Scleral Buckle Implant: Yes (%) | | 12 (54.5) | 4 (50.0) | p=1 |
| ILM Peeling: | Yes (%) | 3 (13.6) | 0 (0) | p=0.793 |

| Patient Data | | No Reoperation | Reoperated | |
|---|---|---|---|---|
| | Previous (%) | 2 (9.1) | 1 (12.5) | |
| | No (%) | 17 (77.3) | 7 (87.5) | |
| Tamponade | C3F8 (%) | 6 (27.3) | 1 (12.5) | p=0.732 |
| | Silicone Oil (%) | 16 (72.7) | 7 (87.5) | |
| Subsequent Silicone Oil Removal: Yes (% of patients under oil) | | 8 (50.0) | 2 (28.6) | p=0.405 |
