## Supplemental Table S3 for "Targeting Rho Kinase in Proliferative Vitreoretinopathy with Netarsudil: Multimodal Preclinical Evidence and Exploratory Clinical Application"

**Table S3**. Antibodies used for Western blotting.

| Target | Antibody | Company (Cat No.) | Dilution |
|---|---|---|---|
| Phospho-Myosin Light Chain 2 (Ser19) | Rabbit pAb | Cell Signaling Technology, Inc. (3671) | 1:1000 |
| Phospho-PTEN (Ser380) | Rabbit pAb | Cell Signaling Technology, Inc. (3671) | 1:1000 |
| Rabbit IgG (Secondary; HRP conjugated) | Goat pAb | R&D Systems (HAF008) | 1:2000 |
