## Supplemental Figures for "Targeting Rho Kinase in Proliferative Vitreoretinopathy with Netarsudil: Multimodal Preclinical Evidence and Exploratory Clinical Application"

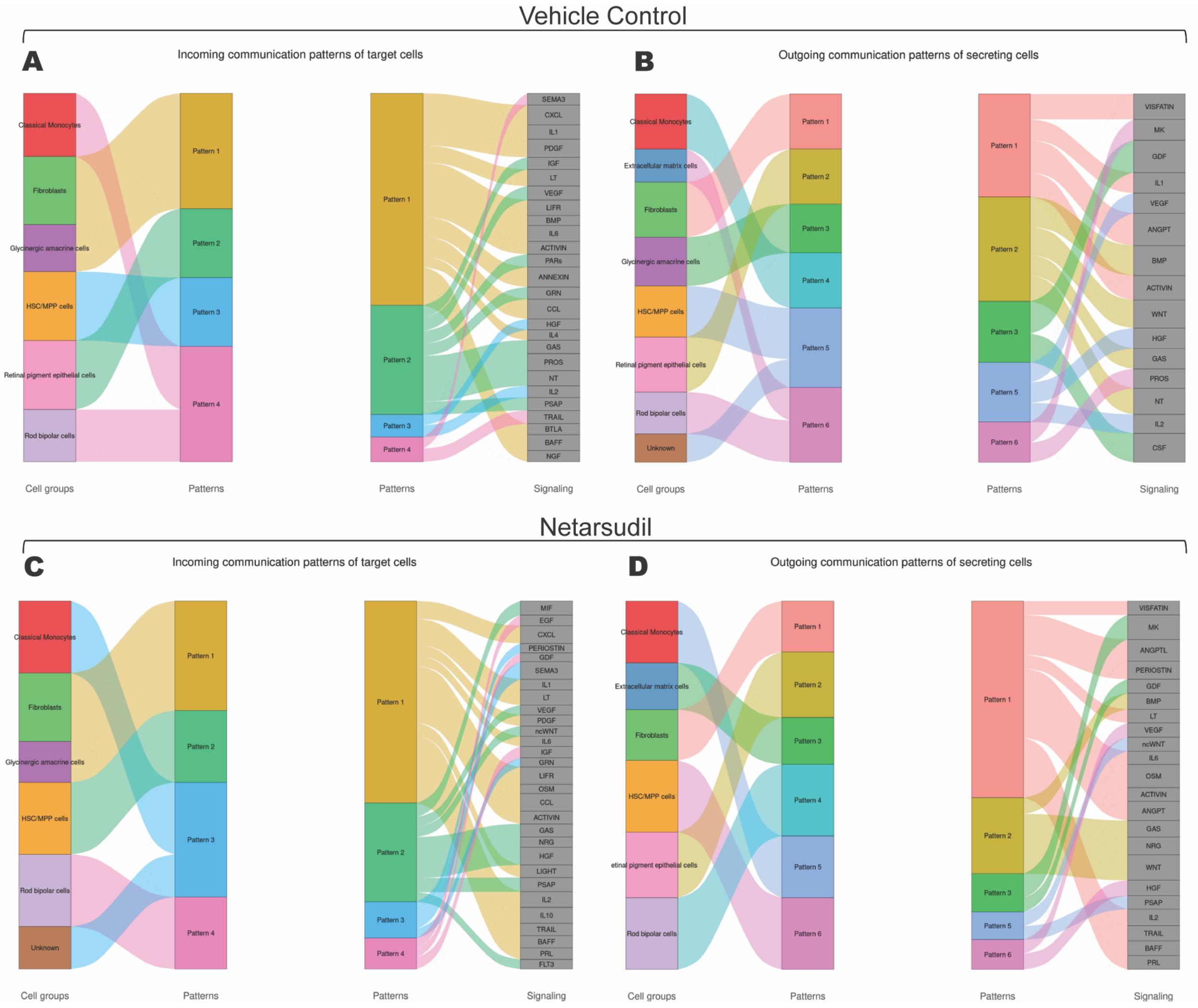


**Figure S1.** Intercellular communication networks inferred using CellChat. **A,B,** Predicted incoming (**A**) and outgoing (**B**) signaling patterns in vehicle control-treated cells. **C,D,** Predicted incoming (**C**) and outgoing (**D**) signaling patterns in netarsudil-treated cells. Differences in communication dynamics highlight the impact of netarsudil on intercellular interactions. The cluster labels shown in this figure are the original scType annotations. After manual review of the gene expression profiles, the annotations were revised to reflect the transcriptional states supported by the data. The original and revised labels are paired below (original → revised). Extracellular matrix cells → ECM component-enriched cells, Rod bipolar cells → Bipolar-like cells, Fibroblasts → Fibroblast-like cells, HSC/MPP cells → Cycling-enriched cells, Retinal pigment epithelial cells → RPE-like cells, Unknown → Unassigned, Classical Monocytes → Monocyte-like cells, and Glycinergic amacrine cells → Neuron-like cells.

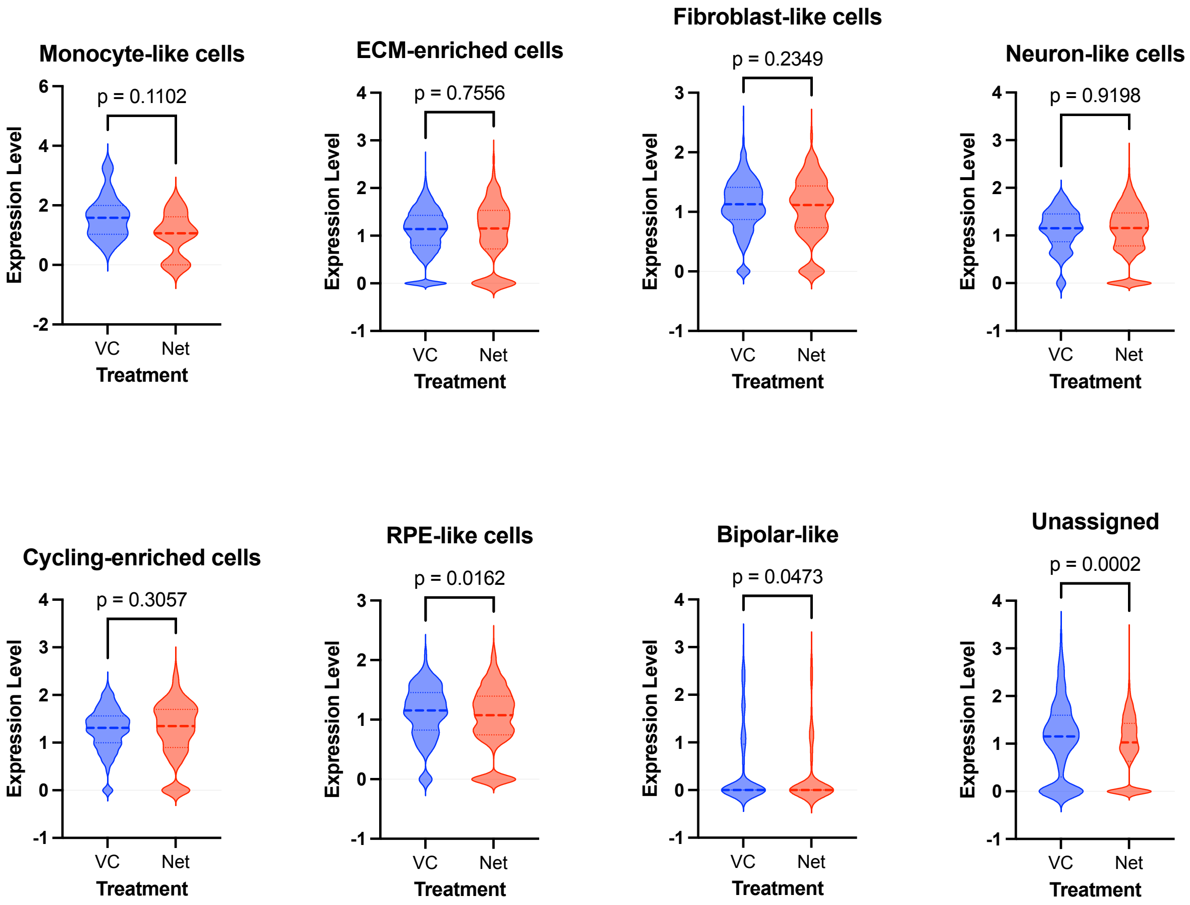


**Figure S2.** RhoA expression across different cell types. RhoA expression levels in (**A**) Classical Monocytes, (**B**) Extracellular Matrix Cells, (**C**) Fibroblasts, (**D**) Glycinergic Amacrine Cells, (**E**) HSC/MPP Cells, (**F**) Retinal Pigment Epithelial Cells, (**G**) Rod Bipolar Cells, and (**H**) Unknown Cells. Differences in expression patterns provide insights into cell-type-specific regulation of RhoA.

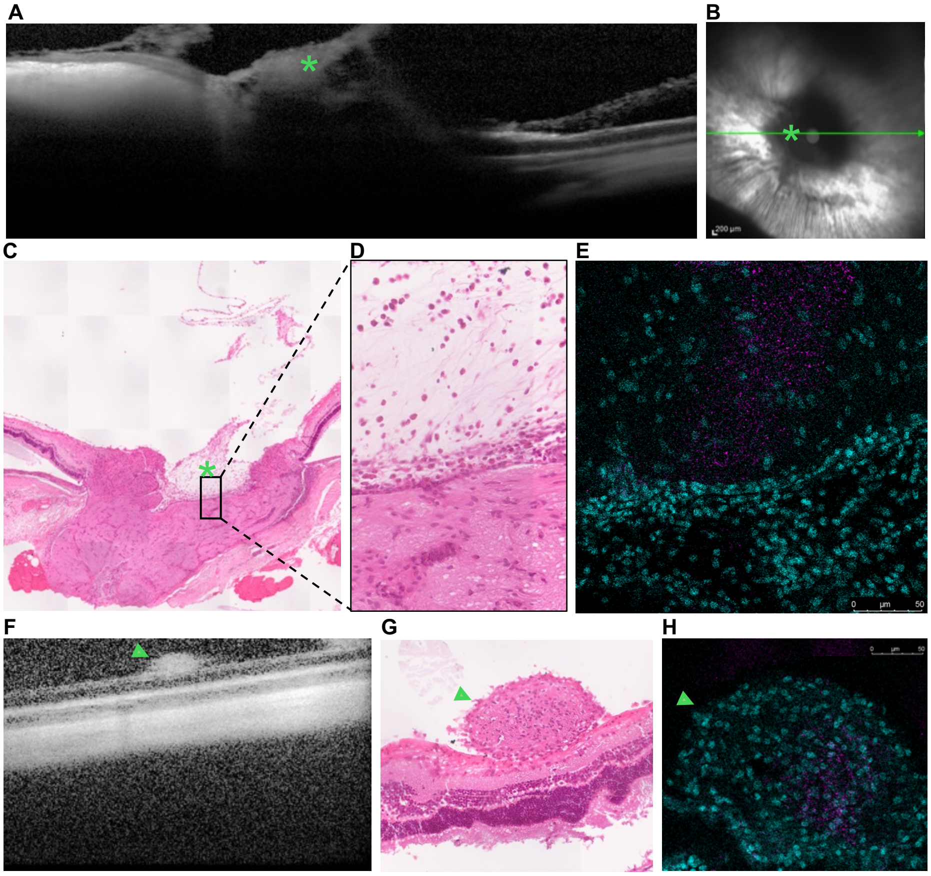


**Figure S3**. Multimodal Indication of residual c-PVR cells in Netarsudil-treated eyes. (**A-E**) Evaluation of the optic nerve head region in a Netarsudil-treated eye, the primary site of c-PVR cell injection. (**A**) OCT scan showing hyperreflective signal at the optic disc (asterisk), consistent with localized pathology. (**B**) Corresponding IR image indicating the scan plane across the optic nerve head. (**C, D**) H&E-stain histologic section of the same region reveals increased cellularity above the optic nerve. (**E**) Immunofluorescence staining with anti-HLA antibodies (majenta) confirms the presence of human c-PVR cells restricted to the the optic nerve head. DAPI (cyan) labels cell nuclei. (**F-H**) Detection of discrete cell cluster on retinal surface in Netarsudil-treated eyes. (**F**) OCT scan revealing a focal hyperreflective round lesion overlying the retinal surface. (**G**) H&E staining demonstrates a cell aggregate independent from the underlying retinal layers. (**H**) Immunofluoresence staining confirms that the cell cluster is composed of human c-PVR cells (anti-HLA, magenta). DAPI(cyan) marks nuclei.
