## Supplemental Methods for "Targeting Rho Kinase in Proliferative Vitreoretinopathy with Netarsudil: Multimodal Preclinical Evidence and Exploratory Clinical Application"

**Supplementary Methods: Single-Cell RNA Sequencing and scType Cluster Annotations**

*Single-Cell RNA Sequencing (scRNA-seq) Data Acquisition and Processing*

Two scRNA-seq datasets were analyzed in this study: one from freshly obtained PVR membranes, which was reanalyzed, and a newly generated dataset from in vitro treated C-PVR cells. The previously published scRNA-seq data was accessed through the Single Cell Portal (Accession: SCP2582; https://singlecell.broadinstitute.org/single_cell/reviewer_access/efb11c05-77ea- 4f1d-8ff8-72461d62190d) [15], and reanalyzed to characterize the transcriptional landscape of the ROCK pathway. The dataset was downloaded as a Seurat object for downstream analysis. The Seurat object was processed using the ScType algorithm for automated cell-type annotation [16,34] as described below. Briefly, a curated gene set from the ScType database was used for classification. The scaled expression matrix was extracted from the Seurat object and used as input for ScType, which assigns cell types based on predefined marker genes. Cell type scores were aggregated at the cluster level, and the highest-scoring cell type was assigned to each cluster. To improve interpretability and reduce redundancy in downstream analysis, ScType-assigned cell types were manually consolidated. Cell types representing cytotoxic and memory T cells as well as natural killer lineages—including Effector $CD8^+$ T cells, Effector $CD4^+$ T cells, γδ-T cells, Memory $CD4^+$ T cells, Natural killer cells, and $CD8^+$ NKT-like cells—were reclassified under the label "Natural Killer/T cells". Similarly, Immature B cells and Plasma B cells were grouped together under the label "B cells", and Starburst amacrine cells were renamed to "Amacrine Cells". This harmonized classification was then used for all subsequent analyses and visualization. Uniform Manifold Approximation and Projection (UMAP) was employed to visualize the annotated cell populations using the DimPlot() function in Seurat (v 5.2.1) [35]. Additionally, ridge plots (RidgePlot()) were generated to display the distribution of selected gene expression profiles across cell types. A custom color palette was applied to enhance the clarity of visualization.

To characterize the molecular effects of netarsudil on C-PVR cells, we performed scRNA-seq on cells treated with either VC or netarsudil (2.5 µM) for 48 hours before dissociation into a single-cell suspension using TrypLE (Gibco) following the manufacturer's instructions. The suspension was sent to the Single-Cell Sequencing Core at Boston University for library preparation and sequencing. Libraries were generated using the 10× Genomics 3' GEM-X single-cell library kit and sequenced on an Illumina NextSeq 2000. Samples from both VC and NET conditions were collected in biological duplicates and processed independently using the 10X Genomics Chromium platform. Sequencing data for each replicate were analyzed separately using the 10X Cell Ranger pipelines for demultiplexing, alignment, cell calling, and unique molecular identifier (UMI) quantification. The cellranger mkfastq command was used to convert raw base call (BCL) files generated by Illumina sequencers into FASTQ format. Alignment and UMI quantification were performed using the cellranger count command with the --expect-cells=10000 parameter.

Droplets with at least 500 UMIs were retained for downstream analysis. These filtered matrices underwent further quality control using the SCTK-QC pipeline [36], which provides a comprehensive suite for detecting low-quality cells, doublets, and ambient RNA contamination. The median number of UMIs per cell was 8,742, and the median number of detected genes was 3,030.

Following individual preprocessing, the count matrices from each VC and NET replicate were merged into a single Seurat object using the Seurat package (v5.2.1) in R (v4.4) [35]. Data normalization and scaling were performed using the SCTransform method, which fits a regularized negative binomial regression model to normalize gene expression values while accounting for sequencing depth and other technical artifacts. Highly variable features were identified using FindVariableFeatures(), and the dataset was scaled to remove unwanted sources of variation. Principal Component Analysis (PCA) was conducted for dimensionality reduction, followed by Uniform Manifold Approximation and Projection (UMAP) for visualization (DimPlot()).

Cell types were classified using the ScType algorithm with curated gene sets specific to eye and immune system tissues [16,34]. Gene sets were obtained from the ScType database and combined into a unified reference for cell type annotation. The scaled expression matrix was extracted from the Seurat object and used as input for ScType, which assigns cell types based on predefined gene sets. Cell type scores were aggregated at the cluster level, and the highest-scoring cell type was assigned to each cluster. To ensure classification accuracy, clusters with scores below a defined threshold (one-fourth of the number of cells in the cluster) were labeled as "Unknown." A custom color palette was applied to enhance visualization clarity.

To compare cell type composition between experimental conditions, the number of cells assigned to each cell type was quantified for VC and NET conditions. These counts were merged into a single data frame for comparative analysis, and normalization was performed by calculating the fraction of each cell type relative to the total number of cells in each condition. Fisher’s exact test was applied to assess significant differences in proportions between VC and NET, and p-values were adjusted for multiple hypothesis testing using False Discovery Rate (FDR) correction with the Benjamini-Hochberg method [https://www.jstor.org/stable/2346101].

Differential gene expression analysis between VC and NET samples was performed using the FindMarkers() function in Seurat. Log-fold change (logFC) thresholds of 0, 0.5, and 1 were applied, and genes with adjusted p-values $< 0.05$ were considered significant. To further explore functional implications, Gene Ontology (GO) enrichment analysis was conducted using the clusterProfiler package (v4.12.6). Genes were mapped to Entrez IDs using biomaRt (v2.60), and enrichment analysis was performed across Biological Process (BP), Cellular Component (CC), and Molecular Function (MF) categories. Enriched GO terms were visualized using bubble plots (ggplot2, v3.5.1), and ridge plots were generated for selected genes.

Gene Set Enrichment Analysis (GSEA) was performed using the fgsea package (v1.30) [37,38]. Ranked gene lists were constructed based on logFC values, and enrichment analysis was conducted using Hallmark pathway gene sets from the Molecular Signatures Database (MSigDB). Pathways with adjusted p-values $< 0.05$ were considered significantly enriched.

Intercellular communication networks were inferred using CellChat (v1.6.1) [39]. For each condition, a CellChat object was created from the Seurat object while retaining cell type annotations. Signaling-related genes were extracted using subsetData(), and analyses were restricted to secreted signaling interactions from the CellChatDB.human database. Overexpressed genes and interactions were identified (identifyOverExpressedGenes(), identifyOverExpressedInteractions()), and ligand-receptor interaction probabilities were computed (computeCommunProb()) with filtering (filterCommunication(min.cells = 10)) to retain interactions present in at least 10 cells. Pathway-specific communication probabilities were refined (computeCommunProbPathway()), and an aggregate network was generated (aggregateNet()). River plots (netAnalysis_river) visualized directional communication patterns between cell types.

**Original scType annotations and revised transcriptional-state labels**

The revised labels reflect the closest transcriptional states supported by manual review of the observed gene expression profiles, without implying definitive cell identity or origin.

| Original scType annotation | Revised label |
|---|---|
| **Extracellular matrix cells** | ECM component-enriched cells |
| **Rod bipolar cells** | Bipolar-like cells |
| **Fibroblasts** | Fibroblast-like cells |
| **HSC/MPP cells** | Cycling-enriched cells |
| **Retinal pigment epithelial cells** | RPE-like cells |
| **Unknown** | Unassigned |
| **Classical Monocytes** | Monocyte-like cells |
| **Glycinergic amacrine cells** | Neuron-like cells |

ECM, extracellular matrix. HSC/MPP, hematopoietic stem cells and multipotent progenitors. RPE, retinal pigment epithelium.
